# CRISPRa screening with real world evidence identifies potassium channels as neuronal entry factors and druggable targets for SARS-CoV-2

**DOI:** 10.1101/2021.07.01.450475

**Authors:** Chengkun Wang, Ravi K. Dinesh, Yuanhao Qu, Arjun Rustagi, Henry Cousins, James Zengel, Tianyi Zhang, Nicholas Magazine, Yinglong Guo, Taryn Hall, Aimee Beck, Lucas Miecho Heilbroner, Grace Peters-Schulze, Aaron Wilk, Luke Tso, Elif Tokar Erdemic, Kae Tanudtanud, Sheng Ren, Kathy Tzy-Hwa Tzeng, Mengdi Wang, Brooke Howitt, Weishan Huang, Jan Carette, Russ Altman, Catherine A. Blish, Le Cong

## Abstract

Although vaccines for severe acute respiratory syndrome coronavirus 2 (SARS-CoV-2) have been successful, there are no good treatments for those who are actively infected. While SARS-CoV-2 primarily infects the respiratory tract, clinical evidence indicates that cells from sensory organs and the brain are also susceptible to infection. While many patients suffer from diverse neurological symptoms, the virus’s neuronal entry remains mysterious. To discover host factors involved in SARS-CoV-2 viral entry, we performed CRISPR activation (CRISPRa) screens targeting all 6000+ human membrane proteins in cells with and without overexpression of ACE2 using Spike-pseudotyped lentiviruses. This unbiased gain-of-function screening identified both novel and previously validated host factors. Notably, newly found host factors have high expression in neuronal and immune cells, including potassium channel KCNA6, protease LGMN, and MHC-II component HLA-DPB1. We validated these factors using replication-competent SARS-CoV-2 infection assays. Notably, the overexpression of KCNA6 led to a marked increase in infection even in cells with undetectable levels of ACE2 expression. Analysis of human olfactory epithelium scRNA-seq data revealed that OLIG2+/TUJ1+ cells--previously identified as sites of infection in COVID-19 autopsy studies-- have high KCNA6 expression and minimal levels of ACE2. The presence of KCNA6 may thus explain sensory/neuronal aspects of COVID-19 symptoms. Further, we demonstrate that FDA-approved compound dalfampridine, an inhibitor of KCNA-family potassium channels, suppresses viral entry in a dosage-dependent manner. Finally, we identified common prescription drugs likely to modulate the top identified host factors, and performed a retrospective analysis of insurance claims of ~8 million patients. This large cohort study revealed a statistically significant association between top drug classes, particularly those targeting potassium channels, and COVID-19 severity. Taken together, the potassium channel KCNA6 facilitates neuronal entry of SARS-CoV-2 and is a promising target for drug repurposing and development.

## Introduction

The emergence of SARS-CoV-2 has led to the COVID-19 pandemic with over 150 million reported cases and over three million reported deaths as of May 2021 (WHO). Coronaviruses are a family of enveloped positive-stranded RNA viruses that cause respiratory and intestinal infections in birds and mammals ^1^. Among the known human coronaviruses, four (229E, HKU1, NL63, and OC43) are widely circulating and cause mild infections, while three (SARS-CoV-1, Middle Eastern Respiratory Syndrome CoV, MERS-CoV, and SARS-CoV-2) are highly pathogenic ^1^.

SARS-CoV-2 enters cells in three major steps. The virus Spike protein first binds to its canonical receptor, angiotensin converting enzyme 2 (ACE2). This is followed by proteolytic processing of the Spike which can be carried out by several proteases, with TMPRSS2 and Furin being the most well-known. These steps lead to membrane fusion and consequent release of viral RNA into the host cell ^2^. Recent studies have also implicated the binding of Spike protein to heparan sulfate ^3^ and cholesterol ^4–6^ as well as soluble-ACE2-mediated host cell attachment ^7^ as factors in viral entry, suggesting additional mechanisms that might be responsible for SARS-CoV-2 tropism and COVID-19 pathology.

Several research groups performed CRISPR loss-of-function (LOF) screens to find factors necessary for SARS-CoV-2 entry and replication ^4,5,8–11^. A large number of hits were only found in one or a few screens but not in others ^12^. Further, several experimentally confirmed entry factors, most notably neuropilins ^13,14^, were not reported as hits in any of the screens ^12^. These discrepancies are likely due to the nature of LOF screens, which can only detect effects of genes that are expressed in the cell lines used.

To date one of the most efficacious treatments of SARS-CoV-2 infection remains antibody-based therapy that targets the SARS-CoV-2 Spike protein to inhibit ACE2 binding and prevent viral entry ^15^. Similarly, current COVID-19 vaccines are highly effective in preventing symptomatic illness and function by triggering an immune response against the Spike protein ^16^. However, newer viral strains with mutated Spike proteins have rendered both antibody therapies and vaccination potentially less effective ^17–21^.

Taken together, this suggests a critical need to gain further insight into SARS-CoV-2 entry mechanisms and develop therapeutics targeting this step in the viral lifecycle. We turned to CRISPR activation (CRISPRa) gain-of-function (GOF) screening of membrane proteins in cells with or without ACE2 receptor using Spike-pseudotyped lentiviruses. Our CRISPRa screen identified hits expressed in a broad spectrum of tissues, revealing previously unknown host factors in neuronal/sensory, respiratory, cardiovascular, and immune systems. We validated some of the most interesting candidate genes—including KCNA6, LGMN, and HLA-DPB1—using cDNA overexpression studies with both pseudoviral and replication-competent SARS-CoV-2 infection assays. Strikingly, we found that exogenous expression of the potassium channel KCNA6 promoted SARS-CoV-2 infection up to ~50 fold in an ACE2-independent fashion. After fixing issues with overlaps in genome annotation, we found that KCNA6 is highly expressed in OLIG2+/TUJ1+ neurons (shown to be infected in COVID-19 autopsy studies despite minimal ACE2/Neuropilin expression) in nasal/olfactory tissues ^13,22^. We then performed a drug-target network analysis and a large retrospective study analyzing insurance claims data, which provided evidence for a clinical association between drug classes targeting our CRISPRa screen hits, especially potassium channels, and COVID-19 severity.

## Results

### Membrane-focused CRISPRa screens identify putative host factors determining susceptibility to SARS-CoV-2

Our CRISPRa screening integrated a dCas9 synergistic activation mediator (SAM) system, a SARS-CoV-2 Spike-D614G pseudoviral entry assay, and a single guide (sgRNA) library targeting all known human membrane proteins (**Fig. 1a, Supplementary Fig.1a**). Specifically, to discover host factors that promote ACE2-dependent and -independent entry of SARS-CoV-2, we engineered two HEK293FT cell lines with or without ACE2 expression (ACE2-null and ACE2-positive) (**Supplementary Fig.1b**). We then engineered them to express the SAM system (**Fig. 1a, Supplementary Fig.1a**) and confirmed the gene activation efficiencies of the cell lines (**Supplementary Fig.1c**). We next generated a pseudovirus-based SARS-CoV-2 entry assay as our functional read-out using a Spike-pseudotyped lentivirus carrying a Zeocin resistance marker fused to a GFP reporter, and confirmed ACE2-dependent pseudoviral infection (**Supplementary Fig.1d**). Of note, we chose to use SARS-CoV-2 Spike protein with the D614G mutation to address the dominance of D614G in global circulating viral sequences. We generated a customized CRISPRa sgRNA library targeting all known human membrane proteins (~6,213 genes totaling ~24,000 sgRNAs with non-targeting controls), and made sure to include predicted membrane proteins that are sometimes absent in genome-wide libraries but could be hidden host factors. We then performed the screening workflow using pseudovirus carrying either SARS-CoV-2 Spike protein (SARS-CoV-2 group) or vesicular stomatitis virus envelope G protein (VSVG group, as a reference) (**Fig. 1a**).

**Fig. 1.**
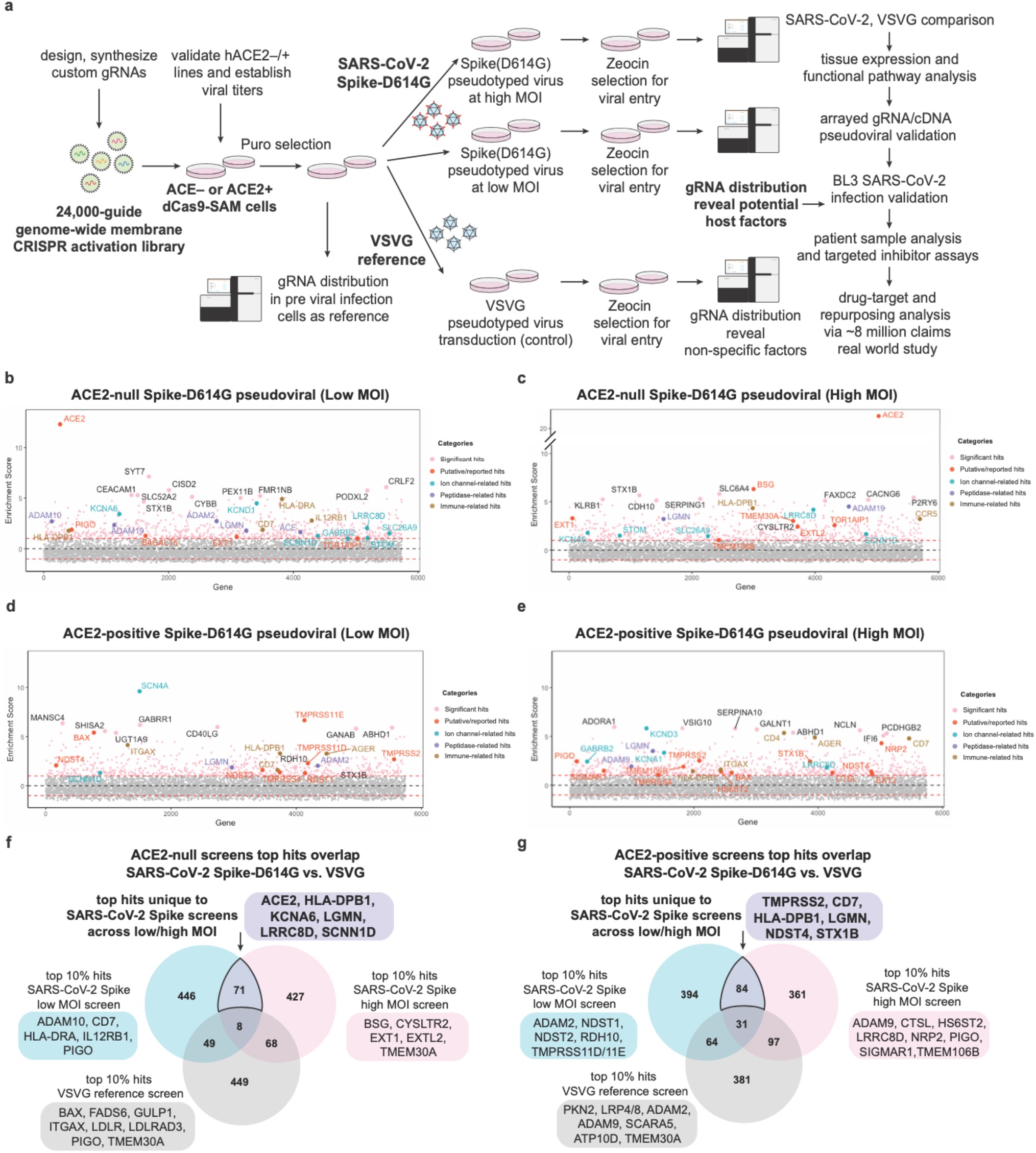
Membrane-focused CRISPRa screening identifies potential host factors involved in Spike-dependent SARS-CoV-2 virus entry. **a,** Screen pipeline showing different conditions used (ACE2-null, ACE2-positive, at low or high MOI, with VSVG references), downstream analyses and validation workflow. **b-e,** Enrichment scores of CRISPRa screen across different conditions with top hits highlighted and colored by their functional categories. **f-g,** Differential analysis of top 10% hits from SARS-CoV-2 Spike and reference VSVG screens. The unique hits in SARS-CoV-2 screens identify putative virus-specific host factors.

We performed the full CRISPRa screen across four conditions: ACE2-null or ACE2-positive at either high or low multiplicity of infection (MOI). We generated at least two biological replicates for each condition and integrated replicates into gene enrichment summaries (**Fig. 1b-e**). The results demonstrated that our approach could identify factors known to promote Spike-dependent viral entry (e.g., ACE2, TMPRSS2, Neuropilin) as well as new genes and pathways (e.g., ion channels, immune and neuronal receptors). First, several genes in the ACE2-null group had significant enrichment that implicated ACE2-independent mechanisms of viral entry (**Fig. 1b-c, Supplementary Fig.2**). Among these genes, some were within known pathways, such as heparan sulfate synthesis enzymes EXT1 and EXTL2 ^3^. Besides known host factors, several ion channel, protease, and immune genes were consistently ranked as top hits, e.g. KCNA6 (a potassium channel), LGMN (a lysosomal/endosomal protease), and HLA-DPB1 (MHC class II beta chain) (**Figure 1f-g**). Second, for the ACE2-positive groups, our screens identified known host factors, including TMPRSS family and NDST enzymes, as well as the recently discovered ACE2-potentiating factor Neuropilin (**Fig. 1d-e, Supplementary Fig.2**). NRP2, in particular, was strongly enriched in the ACE2-positive group. Additionally, several ion channels and neuronal receptors (sodium and potassium channels, GABA receptors) were enriched in this group. A few immune-related genes, such as CD7 and HLA-DPB1, were also among top hits in the presence of ACE2.

### Analyses of top-ranked screen hits identify host pathways involved in SARS-CoV-2 infection and reveal a prominent role for potassium channel genes in viral entry

To examine the top-ranked screen hits, we conducted three analyses: (1) differential analysis compared with VSVG reference screens and previous CRISPR loss-of-function screening studies; (2) tissue expression analysis; and (3) functional pathway enrichment analysis.

First, we sought to identify host factors that specifically promoted SARS-CoV-2 Spike-dependent entry. We ran parallel reference screens using VSVG-pseudotyped lentivirus. As expected, we identified robust enrichment of low-density lipoprotein receptor (LDLR) family members--the canonical VSV host receptors ^23^--among the top hits in the VSVG groups, including LDLR, LDLRAD3, and LRP4/8 (**Fig. 1f-g**). In contrast, known SARS-CoV-2-specific host factors, such as ACE2 and TMPRSS2, were unique to the SARS-CoV-2 screens (**Fig. 1f-g**). Critically, the VSVG reference allowed us to rule out highly enriched genes in SARS-CoV-2 Spike screens that were non-specific. These genes included known non-specific factors involved in apoptosis and growth, such as BAX/PKN2, fatty acid biosynthesis genes PIGO/PIGP/FADS6, and endocytosis/exocytosis genes GULP1/SCARA5. Other potentially pan-lentiviral or pan-viral factors were the integrin ITGAX, and two genes of the P4-ATPase Flippase Complex (the alpha subunit ATP10D, and the beta subunit TMEM30A). In sum, the use of a VSVG reference allowed us to narrow down the range of candidate factors and determine top hits that are specific to SARS-CoV-2 Spike (**Fig. 1f-g**).

In addition, we compare our CRISPRa screens with six recently published CRISPR LOF screens. We extracted the relative rankings of 4923 shared membrane proteins, and generated a curated gene list of 18 validated virus entry factors reported by published studies. We examined the ability of each CRISPR screen to identify these virus entry factors (**Supplementary Fig.3A**). We found that 12 out of the 18 reported entry factors were among the top 10% of the highest ranked genes from our screens (**Supplementary Fig.3**).

Second, we examined the expression of top-ranked genes across 24 tissues using the Genotype-Tissue Expression (GTEx) dataset ^24^ (**Fig. 2a**). Several hits—including STOM, LGMN, and TSPAN15—were broadly expressed across tissues while others showed tissuespecific expression. For example, TMPRSS family genes and SLC26A9 were enriched in the respiratory tract (esophagus and lung), the primary site of SARS-CoV-2 infection. We found that many hits showed high expression levels in neuronal, cardiovascular, liver, and gastrointestinal systems, four organs that have been shown to be involved in the pathophysiology of SARS-CoV-2 ^25–27^ (**Fig. 2b**). Notably, a significant number of our highly-ranked genes are ion channels, transporters or receptors expressed in the brain and sensory systems, including ASIC1, EPHA4, GABRB2, KCNA6, LRRC8D, RDH10, and SCNN1D. Of note, KCNA6 is ranked consistently at top in our screens, but is absent in the latest GTEx database due to overlapping annotations, so we had to study its expression using an alternative source ^28^. Further, we identified liver-enriched hits (e.g. CLEC4G, SIGMAR1) and cardiovascular-enriched hits (e.g. CYSLTR2 and ISLR). Intriguingly, our top hits included several genes highly expressed in immune cells, such as CD7 and MHC-II components. Although there is no direct evidence that immune cells are susceptible to SARS-CoV-2 infection, prior studies have detected SARS-CoV-2 RNA in immune cell types ^29^. Our results potentially explained the presence of SARS-CoV-2 in diverse tissues reported to be susceptible to SARS-CoV-2 ^25–27^.

**Fig. 2.**
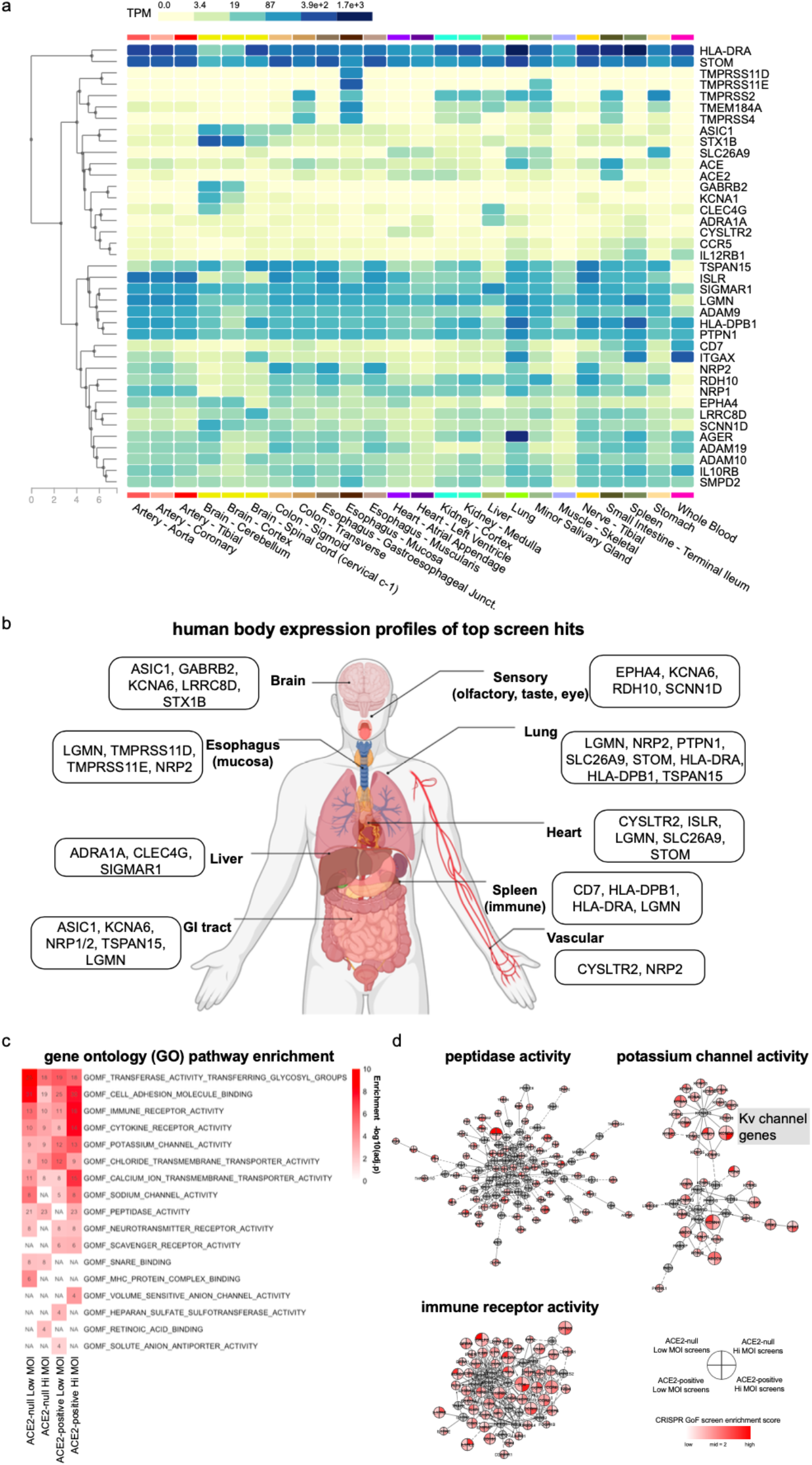
Tissue expression and functional enrichment analysis of top-ranking genes promoting viral entry. **a,** Heatmap showing the overall human tissue expression patterns of top-ranking genes using GTEX v8 dataset. **b,** Tissue expression body map of top-ranking genes, showing putative host genes expressed in the respiratory, neuronal, cardiovascular, liver, gastrointestinal and immune systems. **c,** Gene set overlap analysis using gene ontology (GO) on top 10% of hits from each screen condition. The top GO terms of each screen condition were selected for visualization. **d,** Selected functional network clusters involved with the top GO terms. Colored nodes are significant hits identified from the screen and grey nodes are the connecting nodes. The enrichment score of a gene in each category of screens is indicated by color scale within the node. Notable genes within the same family are highlighted.

Third, we performed functional enrichment and network analyses using top-ranked screen hits. Gene ontology and pathway enrichment (Reactome) analyses identified several biological processes that might relate to SARS-CoV-2 infection (**Fig. 2c**, **Supplementary Fig.4a**). Some were known processes involved in SARS-CoV-2 virus entry, including glycosylation, heparan sulfate synthesis, and peptidase/protease activities ^3,30^. Moreover, some previously unappreciated biological processes ranked highly in the functional analysis, specifically ion channels (potassium, sodium) and immune receptor activities (**Fig. 2c**). To look more closely into these processes, we built detailed functional networks (Reactome FI) using hits within the top-ranked biological processes (**Fig. 2d, Supplementary Fig.4f**) or top screen hits (**Supplementary Fig.4b-e**). One prominent gene family present is the voltage-gated potassium channel (Kv channel) family (**Fig. 2d**). This enrichment of potassium channels and other genes of interest within a functional group pointed to specific host pathways that may mediate or facilitate SARS-CoV-2 entry.

### Pseudoviral and replication-competent SARS-CoV-2 infection validation directly links the expression of top screen hits, most prominently KCNA6, with enhanced viral entry

As part of the initial validation of our screens, we created a subpool CRISPRa library targeting top-ranked genes, transduced them into ACE2-null and ACE2-positive cells, and assayed them with both wild-type (WT) and D614G Spike-pseudotyped lentiviruses (**Supplementary Fig.5**). These experiments confirmed consistent performance of top-ranked sgRNAs irrespective of the use of WT or D614G Spike-pseudotyped lentiviruses (**Supplementary Fig.5**). To validate individual top-ranked genes, we generated focused CRISPRa cell lines in arrayed format and tested them with GFP reporter pseudoviral infection. Activation of high-ranking genes, such as KCNA6, SCNN1D, LGMN, CD7 and MHC-II genes, in ACE2-null cells strongly promoted pseudoviral infection (**Supplementary Fig.6a**). The effects in ACE2-positive cells were less prominent due to ACE2 overexpression, but activation of either LGMN or NRP2 promoted pseudoviral entry at levels comparable to TMPRSS2 (**Supplementary Fig.6b**).

Next, we generated cDNA overexpression cell lines for selected high-ranking genes in ACE2-null and ACE2-positive conditions to directly connect target protein expression with viral entry promotion (**Fig. 3a-b**, **Supplementary Fig.7,8**). We performed arrayed validation on cDNA overexpressing lines using Spike-D614G and VSVG pseudotyped lentivirus. The results of these cDNA experiments differed from the CRISPRa validation, helping to identify *bona fide* entry-promoting effects as unequivocal consequences of exogenous gene expression (**Fig. 3a-b**). In the ACE2-null condition, overexpression of KCNA6 or CD7 specifically promoted Spike-mediated pseudoviral infection, with KCNA6 expression increasing viral entry ~4-fold above the control, compared with ~15-fold for ACE2 cDNA (**Fig. 3a**). In the ACE2-positive condition, overexpression of almost all cDNAs enhanced pseudovirus entry. CD7, EPHA4, LRCC8D, LGMN, and RDH10 overexpressing ACE2-positive lines showed over ~2-fold increases in Spike-mediated infection over the control, and these effects were not seen for VSVG pseudovirus (**Fig. 3b**). Additionally, we used quantitative PCR (qPCR) and Western Blotting to verify cDNA expression in cell lines that promoted Spike-dependent viral entry most robustly (**Supplementary Fig.9**).

**Fig. 3.**
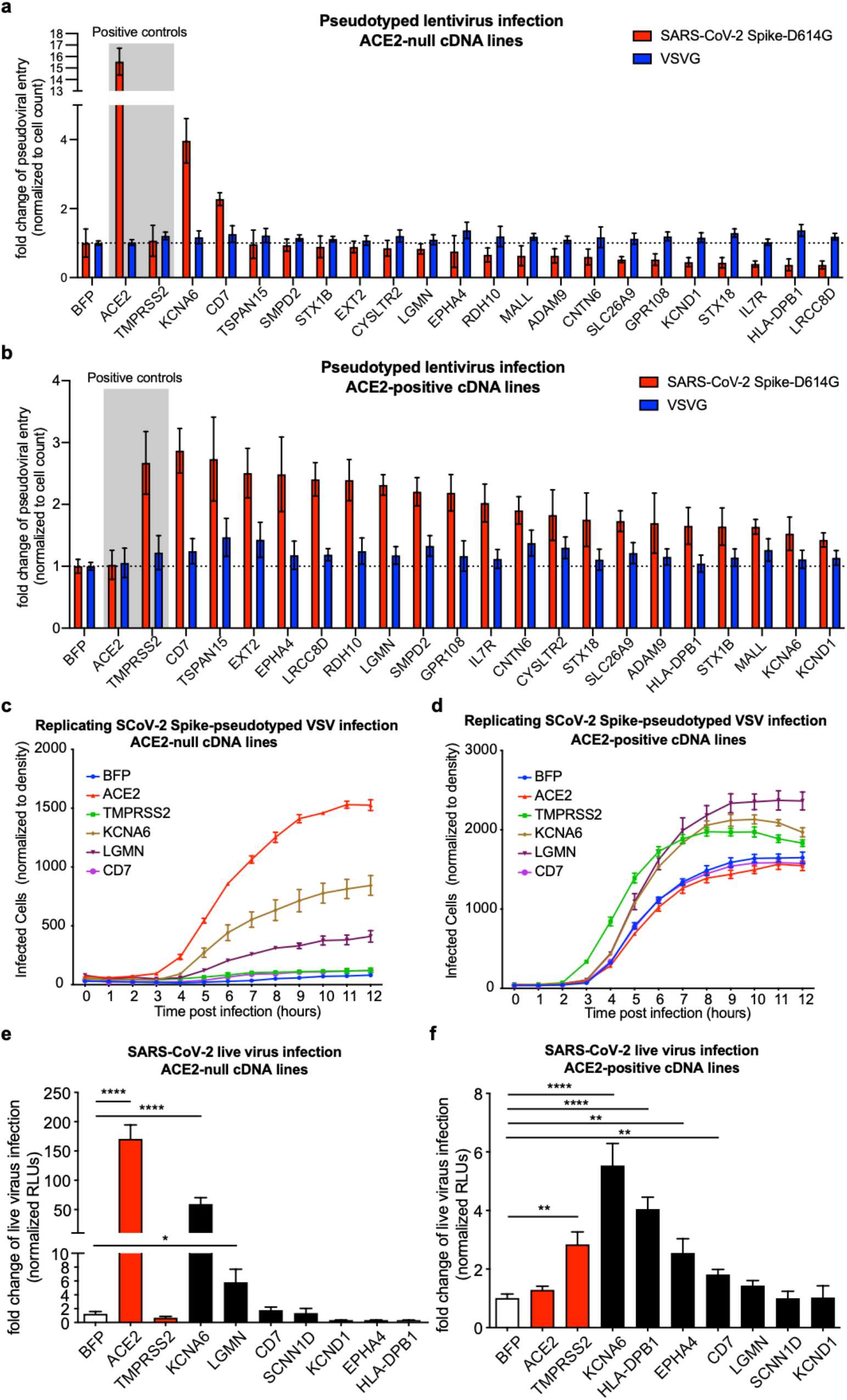
Validation of top-ranked genes using pseudoviral and replication-competent SARS-CoV-2 infection assays. **a-b,** Arrayed validation of top hits in cDNA overexpressing cell lines of individual genes using SARS-CoV-2 Spike-D614G pseudotyped lentiviral assay. The control VSVG-pseudotyped lentivirus results are shown side-by-side. Results are normed to the BFP control for each respective pseudotype. Data from two independent experiments. **c-d,** Arrayed validation using time-lapse imaging of replicating SARS-CoV-2 Spike-pseudotyped VSV infection in cDNA overexpression cell lines. Data from two independent experiments. **e-f,** Validation of top-ranked genes using SARS-CoV-2-nLuc virus infection. Data from three independent experiments. All data represent mean with SEM. Statistical analyses were performed via two-tailed t-test, *, p<0.05; **, p<0.01; ***, p <0.001; ****, p<0.0001.

Using replicating Spike-pseudotyped VSV with time-lapse imaging of virus infection, we found that the strongest hits validated by the pseudotyped lentiviral experiments, such as CD7, KCNA6, and LGMN, were able to similarly promote infection (**Fig. 3c-d**). Parallel experiments with Rabies virus (RABV) G protein pseudotyped VSV in the same cell lines showed no cDNA-dependent effects (**Supplementary Fig.10**). The consistent results from the two pseudoviral systems confirmed that effects from our genes of interest were specific to SARS-CoV-2 Spike, and not artifacts of the systems used.

Finally, we used a nanoluciferase-expressing SARS-CoV-2 reporter virus, icSARS-CoV-2nLuc ^31^, to measure if our candidate overexpression lines could promote live virus infection. Consistent with our pseudovirus results, KCNA6 and LGMN significantly promoted SARS-CoV-2 infection compared with control groups in ACE2-null cells (**Fig. 3e**). Notably, in the ACE2-null condition, KCNA6 overexpression led to a ~50-fold increase in SARS-CoV-2 virus entry, compared with a ~150-fold increase in the ACE2 positive control (**Fig. 3e**). To our knowledge, KCNA6 is the strongest ACE2-independent host factor for SARS-CoV-2 that has been validated by replication-competent virus infection. In addition, KCNA6, HLA-DPB1, EPHA4, CD7, and LGMN also had the ability to promote viral infection in the ACE2-positive condition (**Fig. 3f**). These results demonstrated that our findings were translatable to replication-competent SARS-CoV-2 biology.

### KCNA6 is highly expressed in nasal/olfactory neurons at sites of SARS-CoV-2 infection and pathology

KCNA6 was the strongest candidate host factor from our replication-competent SARS-CoV-2 validation, but it had not been identified in any previous study. We sought to understand why KCNA6 was overlooked and investigate its expression in human tissues to gain insight into its role in viral susceptibility.

As noted earlier, KCNA6 is missing from the GTEx database due to genome annotation overlaps in the latest GRCh38/hg38 genome reference (**Fig. 4a**). Annotations overlapping with KCNA6 are all non-coding: two long non-coding RNAs (RP11-234B24, RP3-377H17, nonconserved in mammals) and a processed non-coding transcript of nearby GALNT8 gene. KCNA6 annotation is also shorter in GRCh38 vs. GRCh37, so that reads may not be effectively mapped to KCNA6 (**Fig. 4a**). Such overlaps led to KCNA6 being entirely discarded from the annotation during preprocessing (the case for GTEx) or having KCNA6 reads mapped to non-KCNA6 annotations. It may thus explain the absence of KCNA6 in prior studies.

**Fig. 4.**
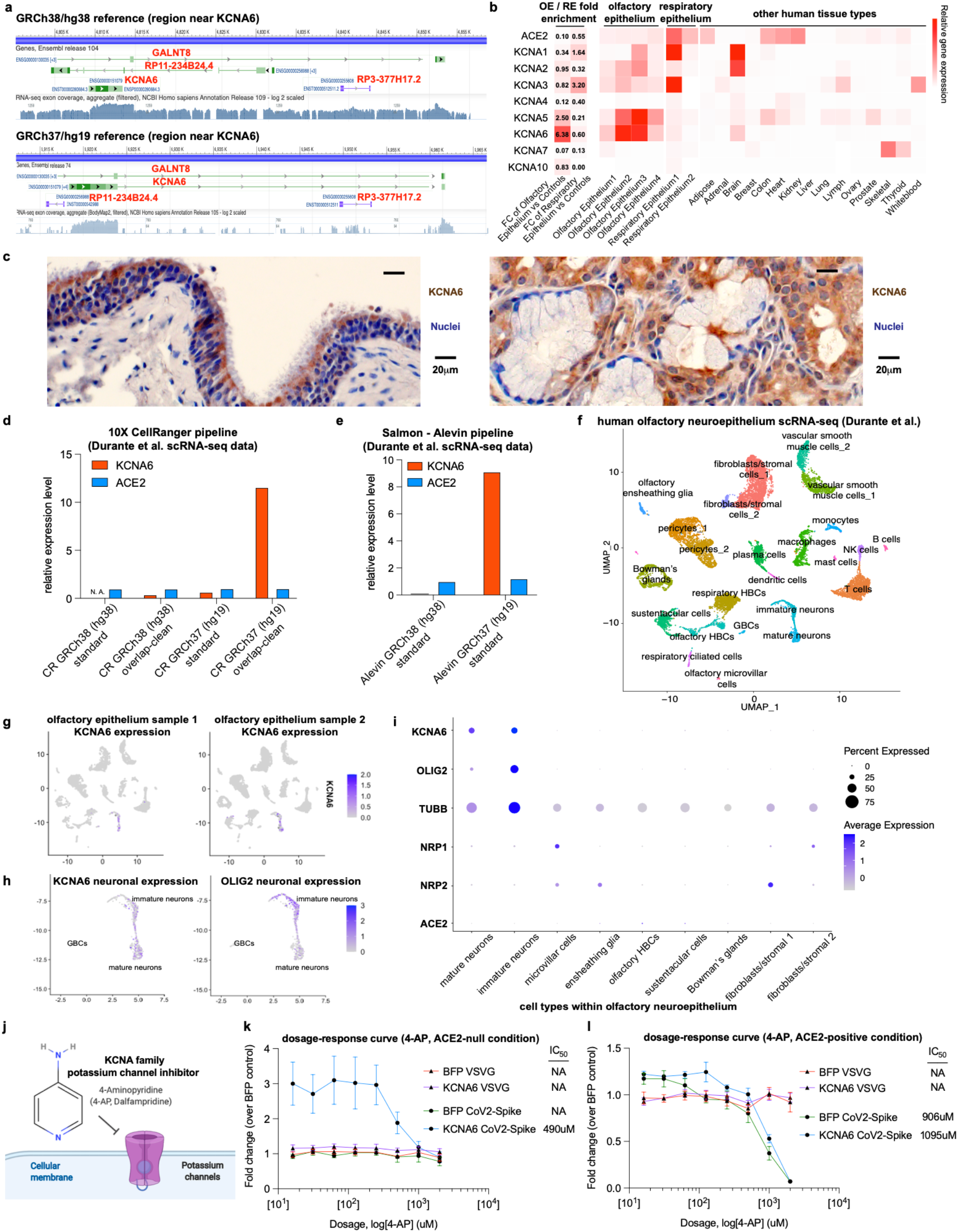
KCNA6 is highly expressed in nasal/olfactory neurons and is a druggable target for inhibiting SARS-CoV-2 viral entry. **a,** KCNA6 genome annotations in the GRCh38 (hg38) and GRCh37 (hg19) references using NCBI Genome Data Viewer. **b,** Expression of ACE2 and KCNA family genes across human tissues. Left two columns show the fold change (FC) of expression comparing olfactory epithelium (OE) or respiratory epithelium (RE) vs. control tissues. Columns on the right show the expression levels in 4 OE samples, 2 RE samples, and 15 control tissues (Olender *et al*., Wang *et al*.). **c**, Immunohistochemistry analysis of KCNA6 protein expression in human sinonasal tissues using archival clinical samples. **d,** Expression of ACE2 and KCNA6 in the single-cell RNA-seq data of olfactory neuroepithelium using different versions of genome references (Durante *et al*.). CellRanger-6.0 was used for all alignments and the expression was calculated by averaging the ACE2/KCNA6 expression in all cells and normalized to the ACE2 expression from the standard GRCh38 reference. **e,** Expression of ACE2/KCNA6 using the Salmon-Alevin pipeline, calculated as in panel **d**. The standard GRCh37/GRCh38 genome references were used. **f,** UMAP depicting the olfactory epithelial cell types from two patients. The cell cluster identities were based on Durante *et al*. **g,** UMAPs depicting the expression levels of KCNA6 in individual patients. **h,** Focused UMAPs of the neuronal populations showing co-expression of KCNA6 and OLIG2, a marker for virus-infected neurons in COVID-19 patient olfactory epithelium.**i,** Dot plot of selected gene expression in olfactory epithelial cell types. Sizes of the dots indicate the proportion of cells having greater-than-zero expression while the color indicates mean expression. **j-l**, FDA-approved compound 4-Aminopyridine (4-AP, dalfampridine) is a broad-spectrum potassium channel inhibitor (**j**). Inhibitor assays performed in ACE2-null (**k**) or ACE2-positive (**l**) conditions, measuring SARS-CoV-2 or VSVG pseudoviral infection efficiencies in KCNA6 cDNA or control BFP lines treated with different doses of 4-AP, with measured IC_50_ to the right. Results are normed to the BFP control/vehicle for each respective pseudotype. Data from two independent experiments and represent mean with SEM.

To understand the true expression of KCNA6 in human tissues, we analyzed two bulk RNA-Seq dataset ^32,33^ using the older genome reference GRCh37/hg19. In the olfactory epithelium, we found that KCNA6 is highly expressed while the canonical SARS-CoV-2 receptor ACE2 is nearly absent (**Fig. 4b**). In the respiratory epithelium, we also detected KCNA6 expression, whereas other KCNA family members and the ACE2 receptor have comparable or higher levels of expression. KCNA6 is mostly enriched in nasal tissues and the brain compared to others (**Fig. 4b**). Further, we performed KCNA6 immunohistochemistry on archival clinical samples of human sinonasal tissues using a previously validated KCNA6-specific antibody (**Supplementary Fig.9**). We observed KCNA6-positive cells with epithelial or serous gland origin. The staining results further verified that KCNA6 protein is present in human nasal tissues at significant levels (**Fig. 4c**).

We then looked into KCNA6 expression in the most enriched olfactory epithelium, taking advantage of a recent single-cell RNA-seq (scRNA-seq) analysis of olfactory neuroepithelium in adult humans ^34^. Given commonly used scRNA-seq pipelines—such as 10x CellRanger, STARsolo, and Kallisto—discard multiple-mapped reads, we found that KCNA6 expression was nearly undetectable when using the latest genome reference GRCh38/hg38 (**Fig. 4d-e**). To fix this issue, we re-processed this scRNA-seq data after adjusting the genome reference to remove overlapping transcripts. Consistent with the bulk RNA-seq results, we now detected high KCNA6 expression using the CellRanger pipeline (**Fig. 4d**). The Salmon-Alevin pipeline can handle multiple-mapped reads, and allows us to readily detect high levels of KCNA6 expression with the standard GRCh37 reference (**Fig. 4e**). KCNA6 is robustly expressed in neuronal cells within human olfactory neuroepithelium (**Fig. 4f-g**). Specifically, we detected high expression of KCNA6 in OLIG2+ neurons (**Fig. 4h**).

Notably, neurons in COVID-19 patient nasal tissues were reportedly infected and involved in disease pathology, but these cells have low expression of ACE2, Neuropilins, or other known entry factors ^35–37^. Two recent studies using COVID-19 autopsy samples have demonstrated that SARS-CoV-2 is detected in OLIG2+ or TUBB(TUJ1)+ neuronal cells of olfactory epithelium ^13,22^. Our analysis showed KCNA6 is uniquely present in this virus-infected cell type (**Fig. 4i**), which can only be seen with updated genome reference. Our findings suggest the presence of KCNA6 could help explain sensory, neuronal aspects of COVID-19 symptoms and suggest the potential for uncovering new SARS-CoV-2 entry mechanisms ^38^.

### Potassium channel inhibition using an FDA-approved compound suggests KCNA6 is a drug target for SARS-CoV-2

Potassium channels are druggable targets. We tested an FDA-approved potassium channel blocker, 4-Aminopyridine (4-AP, dalfampridine), for suppressing viral entry (**Fig. 4j**). The compound 4-AP broadly inhibits potassium channels, most prominently targeting the KCNA-family genes ^39^. Our experiment showed that 4-AP inhibited SARS-CoV-2 pseudovirus entry in a dosage-dependent manner (**Fig. 4k-l**). In ACE2-null groups, 4-AP blocked Spike-mediated viral entry due to KCNA6 expression, with an IC_50_ of ~490uM (**Fig. 4k**). This effect is specific to SARS-CoV-2 as 4-AP had no effect on VSVG pseudoviral infection (**Fig. 4k**). In ACE2-positive groups, 4-AP suppressed the SARS-CoV-2 pseudovirus entry in both the control and KCNA6 lines, but again had no effect on VSVG pseudoviral infection (**Fig. 4l**). In the ACE2-positive group, we observed that KCNA6 overexpression lessened the inhibitory effects of 4-AP (IC50 of 906uM in control vs. 1095uM in KCNA6 cDNA line) (**Fig. 4l**). Here, the modest effect size was likely due to the lower levels of KCNA6 protein in ACE2-positive vs. ACE2-null cells (**Supplementary Fig.9**). Overall, these results suggest targeting potassium channels, particularly KCNA6, as a promising avenue of COVID-19 treatment independent of ACE2-mediated pathways.

### The protease LGMN and MHC-II gene HLA-DPB1 are potential host factors for SARS-CoV-2 infection

From our pseudoviral and replication-competent SARS-CoV2 validation, overexpression of the protease LGMN and the major histocompatibility complex (MHC) Class II component HLA-DPB1, increased viral entry significantly (**Fig. 3b,f**). LGMN, which encodes human legumain/delta-secretase, is thought to be a lysosomal/endosomal asparaginyl endopeptidase (AEP) with broad tissue distribution, and may be activated in an age-dependent manner ^40^. HLA-DPB1 is a human leukocyte antigen (HLA) class II beta chain protein expressed in many human tissues, including the lung. Thus, we examined their single-cell gene expression profiles from published lung bronchoalveolar lavage fluid (BALF) samples of COVID-19 patients ^29^. Our meta-analysis indicated that both LGMN and HLA-DPB1 expression had a positive correlation with levels of SARS-CoV-2 viral RNA (**Supplementary Fig.11**). Moreover, we observed that LGMN expression was positively correlated with COVID-19 disease severity (**Supplementary Fig.11c**). Taken together, LGMN and HLA-DPB1 could be potentially involved in host cell susceptibility to SARS-CoV-2 and COVID-19 disease progression.

### Retrospective analysis of patient claims provides real-world evidence that common drugs targeting potassium channels may lower risks of COVID-19 hospitalization

As our screen targeted all human membrane proteins, the majority of our hits are within the druggable genome. We next asked whether existing drugs are likely to modulate host factors identified in our CRISPRa screens. We constructed a bipartite graph representing known interactions between 4,929 FDA-approved compounds and 2,325 protein-coding genes, 254 of which we identified as SARS-CoV-2-specific screen hits (**Fig. 5a**). To evaluate which drugs were likely to modulate viral entry, we ranked compounds by normalized degree centrality (NDC) with respect to screen hit genes, and identified enriched drug categories by median NDC. We observed marked enrichment of several drug classes with a propensity for potassium-channel targeting, including antidepressant, anticonvulsant, and antipsychotic agents (**Fig. 5b, left**). Many of these classes have previous literature support for a role in modulating SARS-CoV-2 viral entry ^41,42^. To assess the specificity of the highest-ranking drug classes relative to their entire interaction profile, we further scored them by the proportion of drug interactors that were CRISPRa screen hits (degree ratio). This approach identified a similar set of ion-channel-targeting drug categories (**Fig. 5b, right**).

**Fig. 5.**
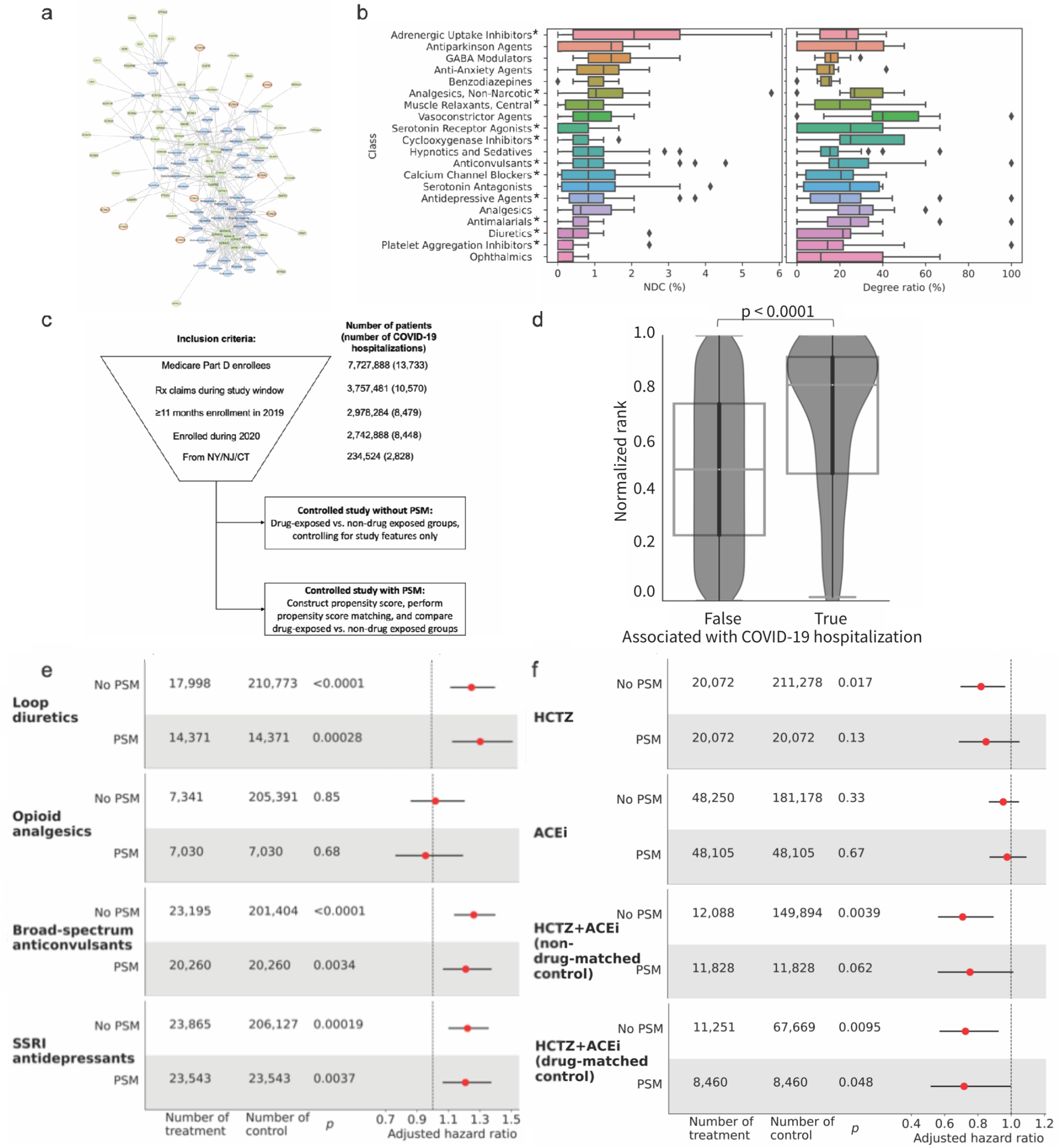
Network analysis identifies drugs that target top screen hits, and retrospective analysis of patient claims provides real world evidence for enriched drug categories. **a,** Overview of the drug-target interaction network, showing an induced subgraph of the 50 highest ranked compounds (drugs in blue; screen hits in green; potassium channel genes outlined in red). **b,** Top drug classes enriched in hits from the interaction network model by NDC and degree ratio with respect to screen hits. Asterisks indicate drug classes with at least one member targeting a potassium channel. **c,** Controlled study design for COVID-19 hospitalization from pharmaceutical claims data. **d,** Drugs associated with COVID-19 hospitalization in the unmatched study rank highly in the drug-target interaction network. **e-f,** Real world evidence for associations between ion-channel-targeting drug classes identified in the screen and increased (**e**) and decreased (**f**) risk of COVID-19 hospitalization in propensity-score-matched subjects.

Next, to provide a clinical assessment of our findings, we performed a retrospective analysis of claims data from a large US health insurance provider, examining associations between common prescriptions and COVID-19 hospitalization rates. We reviewed claims from 7.8 million Medicare Advantage Part D (MAPD) members for compatibility with regional and temporal inclusion criteria (**Fig. 5c**). The final dataset comprised claims for 234,524 MAPD-insured individuals, with at least 11 months of enrollment between January and December 2019 and at least one month of enrollment during 2020, with at least one pharmacy prescription claim from the UnitedHealth Group Clinical Discovery Portal (**Supplementary Table 1**). Among these individuals, 2,828 (1.21%) had claims indicating COVID-19 hospitalization during the observation window.

We screened the claims database for commonly prescribed drugs associated with COVID-19 hospitalization in a 1:10 matched cohort of hospitalized/non-hospitalized patients. We identified 98 drugs whose odds ratio was significant (corrected p<0.05) (**Supplementary Table 2**). This included drugs in several mechanistic classes: broad-spectrum anticonvulsants, angiotensin-converting enzyme (ACE) inhibitors, and thiazide diuretics. These drugs were highly ranked in the drug-target network derived from our screen hits (p<1e^−11^, Mann-Whitney U test; **Fig. 5d**), suggesting an association between a compound’s ability to modulate viral entry genes and its associated risk of COVID-19 hospitalization.

We selected four drug classes for follow-up analysis using 1:1 propensity score matching (PSM). The classes, which included loop diuretics, opioid analgesics, selective serotonin reuptake inhibitor (SSRI) antidepressants, and broad-spectrum anticonvulsants, were ranked highly in the drug-target network, had sufficient prescription volume in the claims dataset, and included known potassium-channel-targeting drugs. The hazard ratios for opioid analgesics were not significant either before or after PSM matching. We observed a significant risk-associated effect for loop diuretics, SSRI antidepressants, and broad-spectrum anticonvulsants that persisted after PSM, with hazard ratios >1.2 consistently and significantly (**Fig. 5e**).

Further, we investigated angiotensin pathway genes, including KCNA channels, which were the most prominent and well-validated category of hits in our experiments. We evaluated whether common drugs targeting these ion channels were associated with decreased risk of hospitalization. Our clinical analysis showed that the diuretic hydrochlorothiazide (HCTZ) alone or in combination with ACE inhibitors (ACEi) were consistently associated with a protective effect, even when compared to controls on other first-line antihypertensive agents, with hazard ratios between 0.72~0.85 and statistical significance (**Fig. 5f**).

## Discussion

In the work presented here we demonstrate the utility of CRISPRa screening to provide insight into the tropism of an emerging pathogen. CRISPR LOF approaches are powerful, but limit discovery of host factors to genes expressed within the context of the cell line used. A potential path to overcome these limitations is doing screens in multiple cell lines representing a variety of cell types, but this is laborious and necessitates prior knowledge of viral tropism. Our CRISPRa screening approach overcomes these limitations by selecting cell lines with limited or no susceptibility and allowing for the unbiased determination of factors that promote viral entry. We show here that previously unknown factors in a diverse set of tissues—such as neuronal (KCNA6), immune (HLA-DPB1, CD7), and cardiac (LGMN, EPHA4)—can stimulate SARS-CoV-2 viral entry.

Most strikingly, we show that KCNA6, a voltage-gated potassium channel, potentiates SARS-CoV-2 entry in a cellular context where ACE2 expression is undetectable (**Fig. 3e, 4h**). Further, our additional tests demonstrated that, under such ACE2-null conditions, KCNA6 overexpression promoted pseudovirus infection when using different variants of the SARS-CoV-2 Spike, including a variant of concern B.1.351 (Beta variant, first identified in South Africa) and B.1.617 (Delta variant, first identified in India) (**Supplementary Fig.12a**). With minimal ACE2 expression, the Spike variants D614G and B.1.351 supported comparable ability as wild-type Spike to infect KCNA6-overexpressing cells in pseudoviral assay (**Supplementary Fig.12b**). This effect maintained when we measured live virus infection using similar set-up. This could be significant in light of reports that emerging SARS-CoV-2 variants could be less dependent on ACE2 binding, and resistant to antibody therapeutics ^17,19,43^.

KCNA6 is highly expressed in nasal OLIG2+/TUJ1+ neuronal cells (**Fig. 4**), which have been shown to be a cell type susceptible to SARS-CoV-2 infection in the olfactory epithelium ^13,22^. Olfactory and taste dysfunction are common and persistent symptoms of COVID-19 ^38^, and a small fraction of hospitalized patients suffer from serious neurological conditions, such as delirium, encephalopathy and stroke ^38,44^. The degree to which these neurological effects are due to infection of neuronal cells or the side effects of an inflammatory state are still poorly understood ^44,45^. Experimental studies have demonstrated that brain organoids are susceptible to SARS-CoV-2 infection despite the low levels of ACE2 receptor expression ^45,46^. Such infection appears to be poorly correlated with levels of ACE2, TMPRSS2, or NRP1 expression ^45^. This suggests that neuronal-specific host factors, such as KCNA6, may work independently or in synergy with Spike binding to ACE2 to promote viral entry.

Prior studies have implicated that host ion channels could promote the entry and replication of several viruses ^47^. In the case of SARS-CoV-2, preliminary work indicated that pharmacological inhibition of pore Ca^2+^ channel 2 (TPC2) led to the decreased entry of SARS-CoV-2 pseudovirions ^48^. A similar finding using MERS pseudovirions found that knockdown of TPC1 or TPC2 lead to decreased viral entry—an effect explainable by lowered Furin cleavage activity and impaired endosomal motility ^49^. Here, we show that FDA-approved KCNA channel blocker 4-AP (dalfampridine) decreased Spike-mediated viral entry (**Fig. 4k-l**). This suggests a potential mechanism that extends beyond KCNA6 to the KCNA potassium channels more generally, which is further supported by our network analysis and real-world evidence (**Fig. 5**). As our work indicates that KCNA6 entry promotion and 4-AP inhibition are both dependent on SARS-CoV-2 Spike, it is interesting to note that the presence of K^+^ ions is a critical requirement for a conformational change in the Bunyavirus envelope glycoprotein that precedes viral entry ^50,51^. Whether a similar mechanism is at play for SARS-CoV-2 Spike is worthy of further investigation.

Our study used SARS-CoV-2 Spike-pseudotyped lentiviruses in our initial CRISPRa screening. As such, our discovery approach does not consider the potential role of other SARS-CoV-2 components, such as the envelope (E) or nucleocapsid (N) proteins. Our focused membrane sgRNA library likely reduced screen noise, as host factors involved in viral entry are rarely non-membrane-associated, but may not detect some potentially interesting proteins that act indirectly to boost infection.

Taken together, the studies presented here offer insight into SARS-CoV-2 viral tropism, yield potential new targets for drug development or drug repurposing, and present a platform that can be applied to future emerging pathogens to understand viral susceptibility.

## Acknowledgments

We are grateful to members of the laboratories of Dr. Le Cong and Dr. Michael Cleary. We are grateful to Dr. Julien Sage for helpful discussion and support on understanding screen results and validation of host factors; to Dr. Joseph Wu, Dr. Mingqiang Wang, Dr. Masataka Nishiga for discussion on single-cell gene expression analysis; to Dr. Weishan Huang and Dr. Tianyi Zhang for helpful discussion on SARS-CoV-2 virus validation.

