## Supplementary Material for "CRISPRa screening with real world evidence identifies potassium channels as neuronal entry factors and druggable targets for SARS-CoV-2"

### Supplementary Information

### Materials and Methods

Plasmids and Constructs

The following constructs were obtained from Addgene: lenti dCAS-VP64_Blast (61425, a kind gift from Feng Zhang), pLX304 (25890, a kind gift from David Root), pCMV-VSVG (8454, a kind gift from Bob Weinberg), pTwist EF1 Alpha nCoV-2019-Spike-2xStrep (141382, a kind gift from Nevan Krogan), pCI-VSVG (1733, a kind gift from Garry Nolan), pXPR_502 (96923, a kind gift from David Root and John Doench), VSV-eGFP-dG vector (31842, a kind gift from Connie Cepko), and pcDNA3.1-hACE2 (145033, a kind gift from Fang Li). EF1a-Hygro, EF1a-ACE2-2A-Hygro, and EF1a-EGFP-2A-ZeoR were Gibson cloned into FUGW using the aforementioned Addgene plasmids as PCR templates. Spike variants (Sd19, Sd19 D614G) were Gibson cloned into the pCI backbone by replacing the VSVG protein in pCI-VSVG. Host factor cDNA containing vectors were ordered from DNASU or Genecopoeia as either lentiviral transfer plasmids or gateway entry vectors. Gateway entry cDNAs were subsequently cloned into the destination vector pLX304 using the Gateway LR clonase kit (Invitrogen 11791019). A list of cDNA vectors is provided in **Supplementary Table 3**.

Cell line culture, generation, and validation

293FT cells were maintained in DMEM with high glucose and Glutamax (Gibco 10566016) supplemented with 1% Pen-Strep (Gibco 15-140-122), 1% NEAA (Gibco 11140050) and 10% FBS (BenchMark) at 37$^{o}$C, 5% $CO_2$.

293FT/dCas9-VP64 cells were generated by transducing 293FT cells with dCAS-VP64_Blast lentivirus. Cells were selected with 10 ug/mL blasticidin and kept on the concentration of selection except in cases of double or triple selection, wherein the doses were reduced to 5 ug/mL. ACE2-null and ACE2-positive cell lines were generated by transduction of either a pLV-EF1a-hACE2-2A-Hygro lentivirus or a pLV-EF1a-Hygro lentivirus into either 293FT cells or 293FT/dCas9-VP64 cells. 293FT or 293FT/dCas9-VP64 cells were selected in 500 ug/mL hygromycin and the doses were reduced to 250 ug/mL in cases of double or triple selection. For cDNA overexpression studies, ACE2-null and ACE2-positive 293FT cells were transduced with cDNA overexpression lentiviruses and selected with blasticidin (10 ug/mL).

Lentiviral, pseudoviral production and transduction

Pseudotyped lentiviruses were produced using PEI or JetOptimus transfection reagent (Polyplus) according to manufacturer's protocols with a ratio of 2:2:1(transfer plasmid: pCMV-dR8.91: Envelope plasmid). For VSVG pseudotyped lentivirus, pCMV-VSVG was used as an envelope plasmid. For SARS-CoV-2 Spike protein pseudotyped lentivirus, pTwist-EF1Alpha-nCoV-2019-Spike-2xStrep was used as an envelope plasmid. For Spike variant pseudotyped lentiviruses, Sd19 and Sd19 D614G variant plasmids cloned in house (as described in Plasmids and Constructs) were used as an envelope plasmid. After 4 hours, the media was replaced and ViralBoost (Alstem, VB100) was added to a 1X concentration. 48 hours after transfection pseudotyped lentiviruses were collected, passed through a 0.45 um filter, and frozen down at -80$^{o}$C.

For large scale production of the membrane protein sgRNA library or production of SARS-CoV-2 Spike protein or Spike variant viruses, 293FT cells in 10 cm dishes were transfected with JetOptimus at the aforementioned ratios, media was replaced 4 hours after transfection, and ViralBoost was added to 1X. At 48 hours, viral supernatant was collected and passed through a 0.45 um filter. For all VSVG psuedotyped viruses, viral supernatant was aliquoted immediately and stored at -80$^{o}$C. For SARS-CoV-2 Spike and Spike variant pseudoviruses, Lentivirus Precipitation Solution (Alstem) was added to 1X. Viral precipitation was carried out according to the manufacturer's protocol. Following precipitation, lentiviruses were concentrated 10X using either DMEM or Ultraculture media (Lonza, discontinued) and frozen down at -80$^{o}$C.

Pseudoviral assay using pseudotyped lentivirus

Every batch of pLV-GLuc-2A-EGFP lentivirus was titered by the addition of 1 x $10^4$ cells to wells of a 96 well plate followed by the addition of varying volumes of virus. Media was added to a final volume of 100 uL and polybrene was added to give a final concentration of 8 ug/mL. Cells were spinfected at 1000 x g, 32$^{o}$C for 45 min before being returned to the 37$^{o}$C incubator. After 24 hours, supernatant was removed from the wells and replaced with 200 uL of fresh media. 72 hours after the spinfection, cells were washed with PBS and resuspended in FACS Buffer (1X PBS, 2% FBS), and assayed for percentages of cells that were GFP positive by flow cytometry. All conditions were done in duplicate. For the experimental assay, 1 x $10^4$ control or modified cells were placed in the wells of the 96 well plate and a volume of virus that gives an MOI of approximately 0.08-0.1 (as determined by the aforementioned titering experiment) was added to each well before the addition of polybrene (8 ug/mL final concentration) and media to a final volume of 100 uL in each well. The assay was then completed as described above for the titering experiment.

Inhibitor Pseudoviral Assay

All inhibitor assays use 96-well plates coated with Poly-D-Lysine (Thermo Fisher, A3890401) at a concentration of 50 ug/mL for 2 hours at room temperature. The plates were then washed with PBS three times, and 1 x $10^4$ cells were plated in a final volume of 100 uL of culture media. The next day, 20 uL of media was removed from each well and replaced with a 5X concentration of the inhibitor in culture media at the indicated dilution. The cells were then returned to 37$^{o}$C. Two hours later, 6 uL of diluted SARS-CoV-2 Spike D614G or VSVG pseudotyped lentiviruses (for an ~MOI of 0.05-0.15) with added polybrene were added to each well for a final concentration of 8 ug/mL polybrene. Plates were spinfected and assayed as described above. 4-Aminopyrimidine (Sigma-Aldrich 36687) was diluted in PBS via vigorously vortexing to a concentration of 100 mM prior to dilution in culture media. The IC50 values were calculated as previously described (Sebaugh, J. L., Pharmaceutical statistics, 2011), which used a four parameter logistic regression model.

Membrane protein CRISPR activation library

A list of all known membrane associated proteins was derived from Chong *et al* (2018), with the following adjustments and manual curation: we included representative olfactory receptor genes and removed pseudogenes or fusion genes that may introduce noise during screening. This refined list was used to pull 4 sgRNAs for each gene from the Calabrese human CRISPR activation pooled library. For genes not included in the Calabrese pool, guides were manually designed using the CRISPOR tool (http://crispor.tefor.net/). For all the final sgRNA sequences, we ensured the starting nucleotide was either G or A by adding a starting G when necessary to maintain efficient Pol-III transcription initiation. The full detail of the customized human membrane protein CRISPRa library design is in Supplementary Data 1.

Oligonucleotide pools were synthesized by TwistBio (CRISPR activation pool) or IDT DNA (additional validation pool) with Esp3I/BsmBI recognition site and PCR amplification sequences appended to the sgRNA sequence. The oligo sequence template was 5'-CATGTTGCCCTGAGGCACAGCGTCTCACACC [guide sequences, 20 or 21 nt] GTTTCAGTCTTCCGTCACATTGGCGCTCGAGA-3'. A set of primers (forward: CATGTTGCCCTGAGGCACAG and reverse: CCGTTAGGTCCCGAAAGGCT) was used to amplify the oligo pool using the manufacturer's protocol (Twist Bioscience, detailed in Twist oligo pool amplification guidelines). The PCR product was column purified using the Monarch PCR and DNA Cleanup Kit (NEB T1030S) and cloned into the pXPR_502 via Golden Gate cloning using the Golden Gate Assembly kit BsmBI-v2 (NEB 1602L). The product was isopropanol precipitated, electroporated into Stbl4 electrocompetent cells (Invitrogen 11635018) with a Micropulser Electroporator (Bio-Rad) in 0.1 cm cuvettes. Cells were allowed to recover in 1 mL of recovery media at 30$^oC$ and then amplified in large scale at 25$^oC$ for 17 hours in 2-YT Broth. Plasmid DNA was prepped using QIAGEN Plasmid Plus Maxi Kit and sequenced to confirm library coverage and distribution.

CRISPRa screening in 293FT cells

For each screen replicate 100 x $10^6$ 293FT/dCas9-VP64 cells with or without ACE2 overexpression were transduced in total in 6 well plates. In each well, 3 x $10^6$ cells were combined with a volume of CRISPRa membrane library viral supernatant to give an MOI of 0.3 before the addition of polybrene (Millipore TR-1003-G) to a working concentration of 8 ug/mL and add culture media to a final volume of 2 mL. Plates were spinfected in a tabletop centrifuge at 1000 x g for 45 min at 32$^oC$. Following spinfection, 2 mL of culture media was added to each well and returned to 37$^oC$ incubators. 12 hours post-spinfection, viral supernatant was removed and cells from each well were split into individual 15 cm dishes at a final volume of 25 mL. 48 hours post-spinfection, puromycin (Gibco A1113803) was added to each plate of transduced and mock-infected cells at a final concentration of 0.5 ug/mL. Cells were selected until mock infected cells were completely killed and transduced cells had recovered to a 90% confluence with minimal cell death in the presence of puromycin selection.

sgRNA containing CRISPRactiv cells were maintained at a minimal cell number of 24 x $10^6$ cells to prevent loss of representation. On the day of the spinfection, 24 x $10^6$ library cells for each condition (ACE2-positive or ACE2-null) were harvested for gDNA. For each screen replicate 100 x $10^6$ library cells for each condition were transduced in total in 6 well plates. For each well, cells were combined with viruses pseudotyped with either SARS-CoV-2 D614G Spike protein or VSVG envelope and carrying the EF1a-BleoR-2A-EGFP transfer vector at an approximate MOI or either 0.01 (low) or 0.1 (high). Polybrene (Millipore TR-1003-G) was added to a working concentration of 8 ug/mL and add culture media to a final volume of 2 mL before cells were spinfected at 1000 x g for 45 min at 32$^oC$. 2 mL of media was added to each well and the plates returned to the 37$^{o}$C incubator until 12 hours post-spinfection when virus containing media was removed and each well was split into individual 15 cm dishes, Zeocin (Gibco R250-01) was added to transduced and mock-infected cells at a final concentration of 500 ug/mL. Cells were selected until mock infected cells were completely killed (approximately 5-7 days post Zeocin addition). Cells for each condition were then pooled and harvested for gDNA extraction.

Genomic DNA isolation, guide RNA amplification and quantification

Genomic DNA of screening samples were extracted using Quick-DNA Midiprep Plus Kit (Zymo Research) following the manufacturer's protocol. Then, 5 ug of genomic DNA was added per 50 uL PCR reaction mixed staggered primers (synthesized by IDT DNA, Forward Primer: 5' AATGATACGGCGACCACCGAGATCTACACTCTTTCCCTACACGACGCTCTTCCGATCT[Stagger, 0-7nt]TTGTGGAAAGGACGAAACACC-3' Reverse Primer: 5'-CAAGCAGAAGACGGCATACGAGAT [8nt-barcode]GTGACTGGAGTTCAGACGTGTGCTCTTCCGATCTTCTACTATTCTTTCCCCTGCACTGT-3') to increase the base diversity. At least 4 PCR reactions were used per sample to ensure adequate coverage. PCR reactions were then pooled and column purified using Monarch PCR and DNA Cleanup Kit (NEB T1030S), visualized on a gel, and column purified following gel extraction. The amplified products were quantified, and normalized by concentration, followed by sequencing using Illumina Miseq Reagent Kit v3 (150 cycles). Saturation analysis was done to confirm sequencing saturation.

Computational analyses of CRISPR activation screens

The sequencing data was deconvoluted using bcl2fastq function (illumina). Reads per each sgRNA were counted and processed using standard Mageck pipeline with an output of RRA and gene ranks. We averaged the positive RRA scores of the biological replicates in each condition and calculated -log (average RRA) scores. Then, enrichment scores were calculated by normalizing -log (average RRA) scores in each condition.

GTEx v8 tissue specific enrichment was performed using the Multi Gene Query function available on the GTEx website: https://www.gtexportal.org/home/multiGeneQueryPage. Gene set overlap analysis was done using top 10% hits in each condition via GSEA-mysigDB (http://www.gsea-msigdb.org/gsea/msigdb/annotate.jsp) with Gene Ontology for Molecular Function. Functional interaction networks were constructed using Reactome FIPlugin in Cytoscape (http://apps.cytoscape.org/apps/reactomefiplugin).

Comparative analyses of SARS-CoV-2 loss of function screens

SARS-CoV-2 loss of function screens data were directly obtained from the supplementary materials of Wei *et al*., Daniloski *et al*., Hoffmann *et al*., Wang *et al*., Zhu *et al*. and Baggen *et al*. on March 31st 2021. Relative rankings of 4923 membrane genes were extracted from each screen. For screens with multiple conditions, the best ranking across the conditions was used. A hit is defined as top 10% if the ranking is equal or smaller than 492.

Generation of validation targeted activation cell lines

Four sgRNAs were designed for every gene of interest and cloned into pXPR_502. sgRNA vectors for a given gene were pooled and these pooled vectors were used to make pooled VSVG pseudotyped lentiviruses using the aforementioned protocol. CRISPR-activated cells were then spinfected at 1000 x g, 32$^{o}$C, for 45 min in an arrayed format with the pooled lentiviruses for a given gene in 48 well plates at a density of 5 x $10^4$ cells/well. 48 hours after spinfection**,** cells were split into 6-well plates and selected with 0.5 ug/mL puromycin until they recovered to 70-80% confluence with no indication of continued selection induced cell death. These cells were then subjected to a SARS-CoV-2 spike D614G pseudotyped lentiviral assay as described in the previous section.

Generation of focused validation libraries

For ACE2-null and ACE2-positive screens, we separately selected the top hits that showed up in multiple biological replicates and MOI conditions and curated lists of genes for the focused pooled validation: 523 genes for ACE2-null library and 542 genes for ACE2-positive library. We added 2 guides per gene on top of the original library (6 guides per gene total) and 30 non-targeting guide RNAs as control. The oligonucleotide pools were also synthesized by Twist Bioscience. The plasmid library and lentivirus were made as described in the previous section.

Analysis of RNA-seq and scRNA-seq data

The human olfactory epithelium RNA-seq data were directly obtained from Olender *et al* (https://www.ncbi.nlm.nih.gov/pmc/articles/PMC4982115/) and visualized using pheatmap in R. For scRNA-seq of the olfactory epithelium, patient 2 and patient 3 data (BAM file) from Durante *et al.* were downloaded from NCBI SRA portal (GSE139522). Because of the overlap of the references, we have built a customized version of GRCh37 reference by deleting the references of RP11-234B24.4 and GALNT8 in the GRCh37.87. Then the data were aligned using the customized GRCh37 reference via CellRanger 6.0 (10X Genomics). For Alevin single-cell analysis, standard genome references (GRCh38 and GRCh37) built with Salmon (https://github.com/COMBINE-lab/salmon) were used to process the same datasets as CellRanger. The Salmon - Alevin pipeline was used with default parameters and adjustment of the 10x kit version according to the dataset specifications, namely 10x-v3 for patient2 and 10x-v2 for patient3. The result data matrices were processed using Seurat following the methods mentioned in Durante *et al*. (https://github.com/satijalab/seurat). Bronchoalveolar lavage fluid (BALF) data from Liao *et al* was acquired from https://github.com/zhangzlab/covid_balf and then processed and visualized using Seurat in R.

Flow cytometry analysis and surface marker staining assays

For ACE2 staining, cells were washed with 1X PBS and then incubated with biotinylated SARS-CoV-2 Receptor Binding Protein (RBD) (ACRO Biosystems SPD-C82E9) at a concentration of 4 ug/mL in FACS Buffer (1X PBS with 2% FBS) for 30 min at room temperature. The cells were washed twice with FACS Buffer and then incubated with streptavidin-Alexa 488 (Thermo Fisher, S11223) at a concentration of 2 ug/mL for 30 min at room temperature. Cells were washed twice with FACS Buffer and analysis was carried out on a Cytoflex Flow Cytometer. For CD46 and CD55 staining, cells were washed with 1X PBS and then incubated with either APC anti-human CD46 antibody (Biolegend 352405) or APC anti-human CD55 antibody (Biolegend 311311) diluted 1:100 in FACS Buffer for 30 min at room temperature. Cells were then washed twice with FACS Buffer and analyzed as described above.

Western Blotting

Cells were pelleted, washed with 1X PBS, and lysed in RIPA buffer (Cell Signaling Technology 98306). Immunoblotting was performed with the following primary antibodies: V5 (Thermo R960-25) and KCNA6 (Sigma HPA021516). All procedures and dilutions were performed according to the manufacturer's recommended protocol.

Immunohistochemistry

All tissues were obtained from the Stanford Tissue Bank in compliance with all institutional guidelines. The tissue blocked were sectioned and then the immunohistochemistry was conducted on series of 4-μm-thick formalin-fixed, paraffin-embedded tissue sections using rabbit anti-KCNA6 polyclonal antibody, applying the Vectastain Elite ABC Universal Plus Kit according to their standard protocol. The following antibodies and dilutions were used: KCNA6 polyclonal antibody (HPA014418, Millipore-Sigma), and the dilution factor for all KCNA6 stainings is 1:500. The antigen retrieval for each stain included a 20-minute heat-activated epitope retrieval in pH 6 citrate buffer using a steamer.

Replicating vesicular stomatitis virus (VSV) pseudovirus generation

Recombinant VSV expressing eGFP in the 1st position (VSVdG-GFP-CoV2-S) was generated as previously described (https://doi.org/10.1016/j.cell.2020.12.004). The plasmid to rescue this virus was generated by inserting a codon optimized SARS-CoV2-S based on the Wuhan-Hu-1 isolate (Genbank:MN908947.3), which was mutated to remove a putative ER retention domain (K1269A and H1271A) into a VSV-eGFP-dG vector (Addgene, Plasmid #31842) in frame with the deleted VSV-G. The control virus VSVdG-RABV-G SAD-B19 was also generated by inserting Rabies virus G in the same vector. Both viruses were rescued in 293FT/VeroE6 cell co-culture and amplified in VeroE6 cells and titrated in VeroE6 cells overexpressing TMPRSS2. Sequencing of the amplified virus revealed an early C-terminal Stop signal (1274STOP) and a partial mutation at A372T (~50%) in the ectodomain. Similar adaptive mutations were found in a previously published VSVdG-CoV2-S (https://doi.org/10.1016/j.chom.2020.06.020).

Pseudovirus infection assay using Replicating VSV pseudovirus

HEK293FT cells were plated in clear 96-well plates at $2x10^4$ cells per well approximately 24 hours prior to infection in 100 uL of media containing 10% FBS. Cells were infected with VSVdG-CoV2-S or VSVdG-RABV-G at an MOI of 0.1. Infection was performed by diluting virus in media without FBS and adding 150 uL of diluted virus per well. After addition of virus, the plate was spun at 900 x g for 60 minutes at 30°C. Infection was tracked over time using an Incucyte system (Sartorius) in a 37°C and 5% $CO_2$ incubator using 4x magnification and detecting GFP. GFP+ cells were counted using Incucyte Analysis software and data was reported as GFP positive foci per well after normalization to confluence.

Replication-competent SARS-CoV2 live virus infection assay

SARS-CoV-2-nLuc (https://www.nature.com/articles/s41586-020-2708-8) in the form of a passage 1 stock was a kind gift from Jacob Hou and Ralph Baric. The virus was passaged twice in VeroE6 cells and titered by plaque assay on VeroE6 cells. Cells were plated in solid white 96-well plates. Cells were then infected at MOI ~0.1, washed, and incubated for 48 hours before assessment by lytic Nano-Glo assay (Promega) and read on a GloMax plate reader (Promega). Infections and plate reading occurred inside class II biosafety cabinets under biosafety level 3 (BL3) conditions. All experiments using viruses were approved by the Administrative Panel on Biosafety (APB).

Drug-target network analysis

Genes were labeled as screen hits by first selecting the top quartile of genes by enrichment score, then removing genes that were also hits in the VSVG screen (>95$^{th}$ percentile by enrichment score). A list of drug-gene interactions was obtained from DrugBank and used to generate a bipartite graph with two node classes, drugs and genes, and edges representing known interactions between protein-coding genes and FDA-approved drugs. From this graph, the “full network,” a subgraph was generated containing only genes that were screen hits and their associated drug interactions (the “screen-hits network”). Drugs were labeled according to DrugBank classifications and were ranked by normalized degree centrality as defined below:

$$ndc = n^{screen\ hits} / m^{screen\ hits}$$

where $n^{screen\ hits}$ is the node degree in the screen-hits network and $m$ is its maximum theoretically possible degree in the same network.

Drug nodes were separately ranked by “degree fraction,” as defined below:

$$degree\ fraction = n^{screen\ hits} / n^{full\ network}$$

where $n^{screen\ hits}$ is the node degree in the screen-hits-only network and $n^{screen\ hits}$ is the node degree in the full network.

When calculating aggregate rankings for drug classes, we included only classes with greater than 10 members in the full network. Statistical tests are Mann-Whitney U tests and Spearman rank-order correlation as appropriate, unless stated otherwise.

Claims database

The study sample was obtained from de-identified administrative claims for Medicare Advantage Part D (MAPD) members in a research database from a single large US health insurance provider (the UnitedHealth Group Clinical Discovery Portal). The database contains medical (emergency, inpatient, and outpatient) and pharmacy claims for services submitted for third party reimbursement, available as International Classification of Diseases, Tenth Revision, Clinical Modification (ICD-10-CM), and National Drug Codes (NDC) claims, respectively. These claims are aggregated after completion of care encounters and submission of claims for reimbursement.

Database-wide drug screen

For the initial claims database screen, drugs were defined by their generic names and used by individuals represented in claims data between July 1, 2019, and January 31, 2020. We selected the screening study cohort to be all the COVID-19 related hospitalized members and 1:10 exactly matched non-hospitalized members based on: age (+/- 1), gender, race, socioeconomic status (SES) index (+/- 0.5), living in counties from New York, New Jersey and Connecticut or counties outside the New York, New Jersey and Connecticut tri-state area, and diagnosis of diabetes without chronic complications, congestive heart failure, chronic pulmonary disease, myocardial infarction, metastatic cancer, liver disease, renal failure, peptic ulcer disease, and hypertension. We included unique drugs with over 2,000 users in the 1:10 exact matched cohort in the screening study. For each drug, we considered individuals drug-exposed if they had any reimbursed prescription claims during the study period. We considered individuals non-drug exposed if they had zero reimbursed prescription claims for the analyzed drug during the study period. We calculated the log odds of COVID-19 hospitalization as a binary outcome, comparing drug-exposed to non-drug-exposed individuals, across drug claims meeting our minimum sample size threshold. Odds ratio significance levels were adjusted for multiple hypothesis testing using a Benjamini-Hochberg correction.

Cohort construction

Drug classes were defined by American Hospital Formulary Service (AHFS) codes**.** For each drug class of interest, we constructed a cohort of individuals with at least 11 months of enrollment in MAPD insurance from January through December 2019 and at least 1 month of enrollment in MAPD in 2020. These individuals had at least one pharmacy prescription claim during their enrollment and lived in counties in New York, New Jersey, and Connecticut (**Supplementary Table 1**). In our database, COVID-19 hospitalization is more prevalent among individuals insured through MAPD and among residents of the New York, New Jersey, and Connecticut tri-state area. We restricted our analyses to these populations to select for uniform exposure to COVID-19 and a higher prevalence of the COVID-19 hospitalization outcome in our cohort. We define our outcome as a claim for a hospitalization with a positive COVID-19 test between January 1, 2020 and June 26, 2020.

Prescription drug users were identified by string matching from pharmacy claims for any of the generic names associated with the drug candidate. We considered individuals to be drug-exposed when their total supply days covered ≥80% of days between their first drug use date after July 1, 2019 through January 31, 2020. We considered individuals non-drug-exposed if the individual was never prescribed the drug candidates or drugs in the same therapeutic class, between July 1, 2019 and January 31, 2020. We also included one negative control associated with a known COVID-19 confounder, glucose meters, to assess our analysis pipeline's global confounding control. We considered individuals to be exposed when they have one prescription for a glucose meter between July 1, 2019 and January 31, 2020.

Study covariates

For each drug of interest, we extracted the following list of covariates for both drug-exposed individuals and non-drug-exposed individuals:

1. Age
2. Gender
3. Self-reported race and ethnicity
4. Area-specified SES index based on member zip code
5. 2019 diagnoses as selected from the top 200 first three-digit ICD-10-CM code, excluding codes beginning with “Z”
6. Pre-existing conditions defined by diagnosis codes in 2019, including conditions used in the Charlson Comorbidity Index and Elixhauser Comorbidity Index
7. Pre-existing primary treatment-related diagnosis
8. Co-used prescription drug defined as claims between July 1, 2019, and January 31, 2020, for the top 20 therapeutic classes
9. Prior hospitalizations in 2019
10. Count of primary care provider visit in 2019
11. Count of unique drugs prescribed
12. Routine screening adherence in 2019, as indicated by completion of a comprehensive metabolic panel, lipid panel, and complete blood count
13. Flu vaccination in 2019 as a proxy of good health behaviors
14. Special Need Plan: (1) institutional, indicating if a member is from a nursing home; (2) dual plan with Medicaid.

<u>Controlled study without propensity score matching</u>

We first selected a list of features using a LASSO model with tuned penalty coefficient based on Bayesian information criteria. The complete list of features includes normalized age, sex, primary treatment-related diagnosis, comorbidity index flags, occurrence flags to first three digits of diagnosis codes, adherence flags to co-used drug therapeutic classes, race, state of residence, and normalized SES index. After feature selection, we added normalized age, normalized SES index, and primary treatment-related diagnosis into the feature list to control for these factors. To ensure model convergence, we excluded features with a prevalence of less than one percent of the cohort. We then fit a Cox proportional hazard model to determine the adjusted hazard ratio of the treatment group, considering time to COVID-19 hospitalization, controlling for the list of features selected. We allowed baseline time to vary by individual, setting individual baseline time to be time in our database of first COVID-19 hospitalization for an individual residing in the same state.

<u>Controlled study with propensity score matching</u>

For the group of drug-exposed individuals, we applied 1:1 propensity score matching (PSM) to construct a matched group of non-drug exposed individuals. The propensity score was built using logistic regression based on age, sex, primary treatment-related diagnosis, comorbidity index flags, occurrence flags to first three digits of diagnosis codes, adherence flags to co-used drug therapeutic classes, race, state of residence, and SES index. We ran 1:1 PSM with a caliper of 0.25 multiplied by the standard deviation of propensity scores. We assessed PSM performance by calculating the standardized mean difference between drug-exposed and non-exposed groups across the primary treatment related diagnosis. PSM is considered adequate when the standardized mean difference between groups is ≤ 0.10 (Zhang *et al.*, 2019). After PSM we report the unadjusted hazard ratio for the drug-exposed group. In addition, we applied the same procedure of feature selection and similarly fit a Cox proportional hazards model for each drug of interest, between baseline (the state-specific time of first COVID-19 hospitalization) to hospitalization or end of follow-up, to investigate the adjusted hazard ratio of the drug-exposed group.

Figure panels 1A, 2B are adapted from templates or created with BioRender.com (2020).

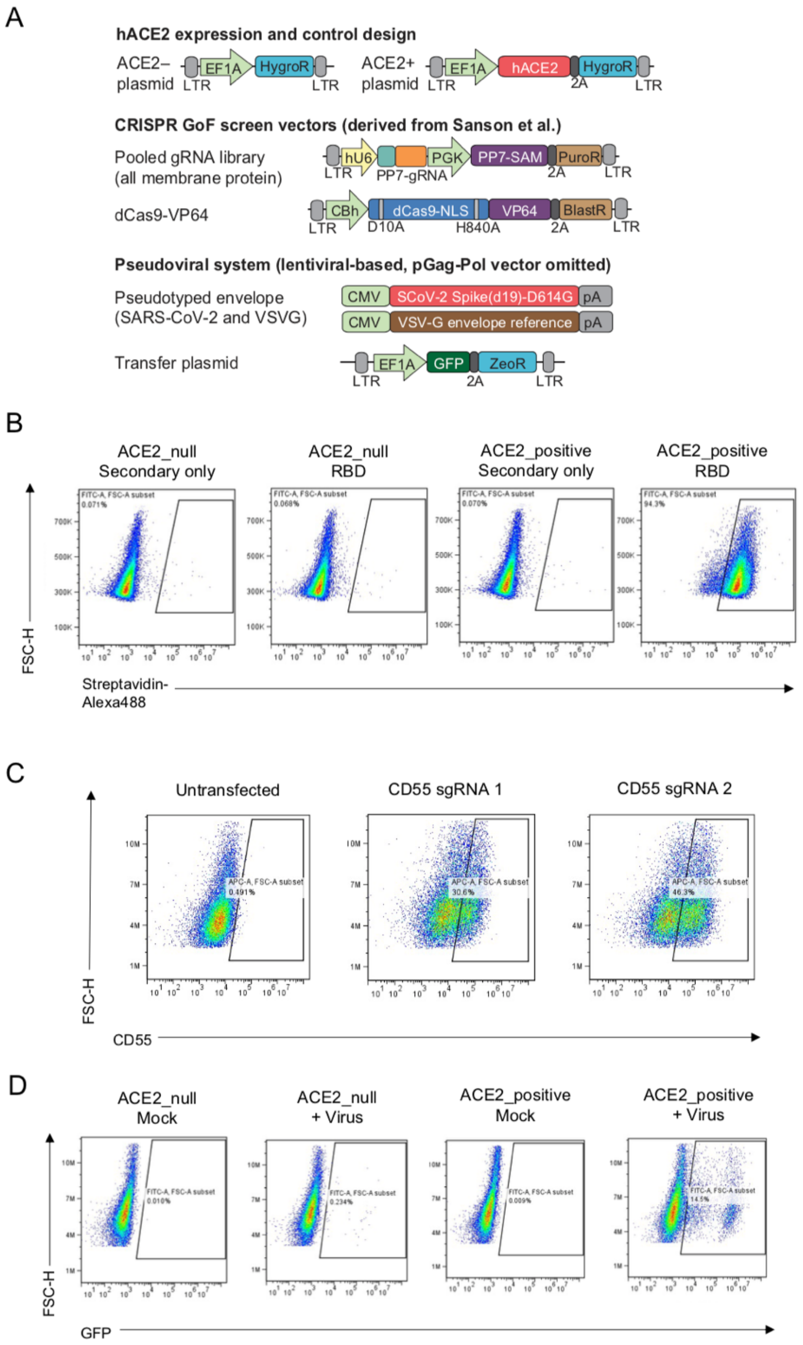


**Supplementary Fig. 1. Development of a pseudoviral based platform to screen for novel SARS-CoV-2 entry factors.** (a) Schematics showing the design of vector systems used in CRISPRa screening. (b) ACE2-null and ACE2-positive lines stained with or without RBD-Biotin and Streptavidin-Alexa488. (c) ACE2-null-dCas9-VP64 cells stained with CD55-APC five days after transfection with pXPR_502 plasmids encoding sgRNAs targeting the promoter of CD55. (d) ACE2-null and ACE2-positive cells either mock infected or infected with SARS-CoV-2 Sd19 pseudotyped lentiviruses.

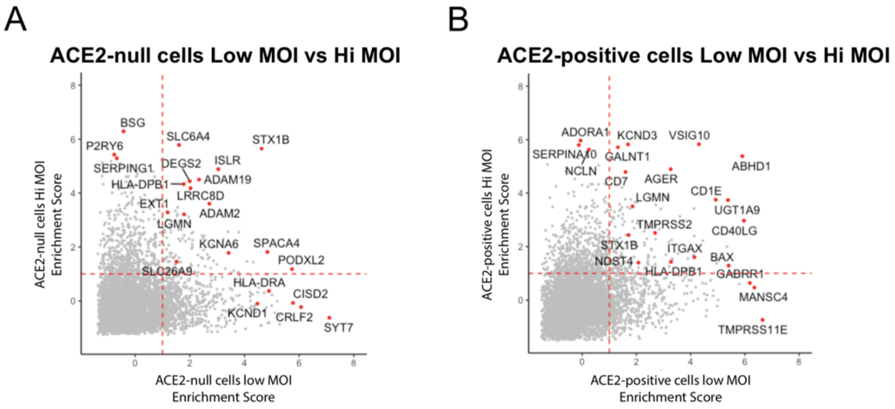


**Supplementary Fig. 2. Comparison of the screen enrichment scores using low (~ 0.01) or high (~0.1) MOI of SARS-CoV-2 614G Spike pseudotyped virus.** (a-b) Scatter plots show the enrichment scores from (a) ACE2-null 293FT cells or (b) ACE2-positive 293FT cells.

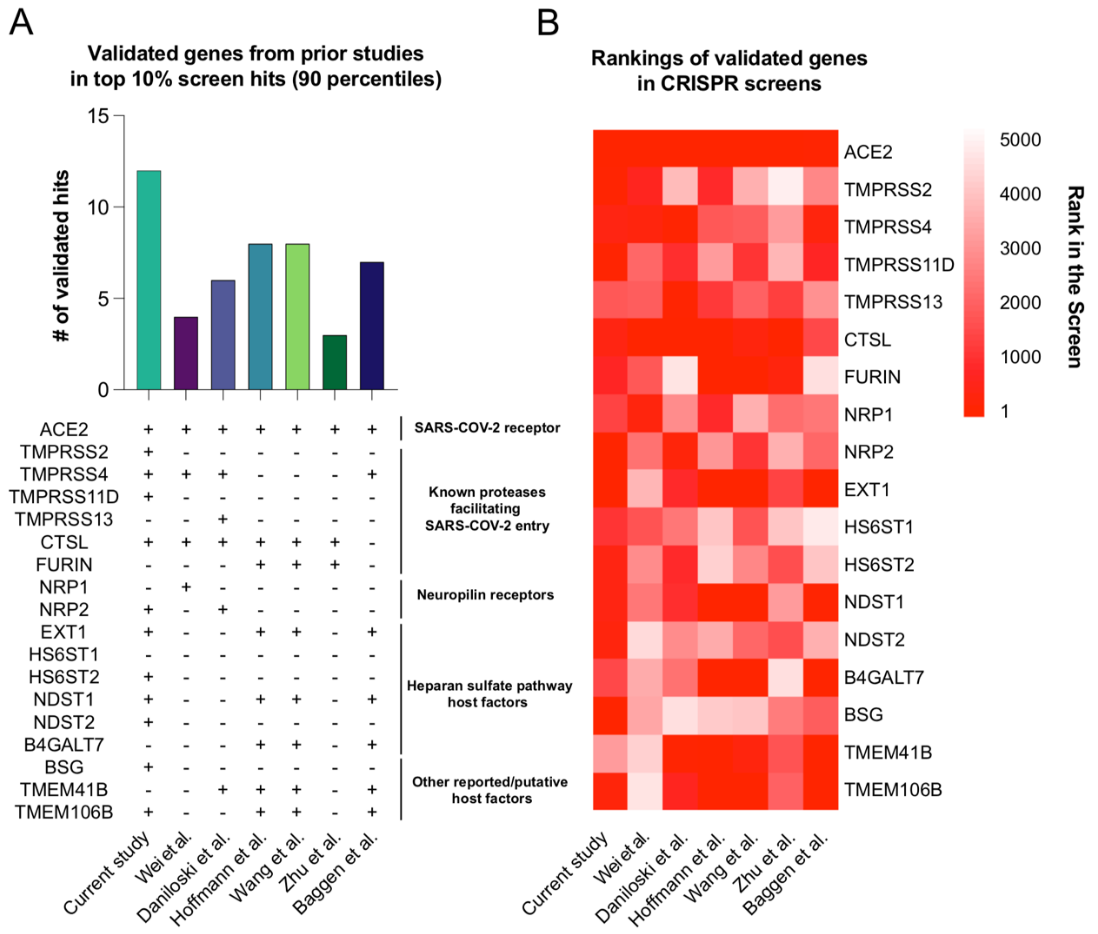


**Supplementary Fig. 3. Identification of the established membrane entry factors for SARS-CoV-2 in our screens and previous loss-of-function screens.** (a) Bar plot shows the number of established membrane entry factors in the top 10% screen hits (b) Heatmap shows the relative ranking of the established membrane entry factors.

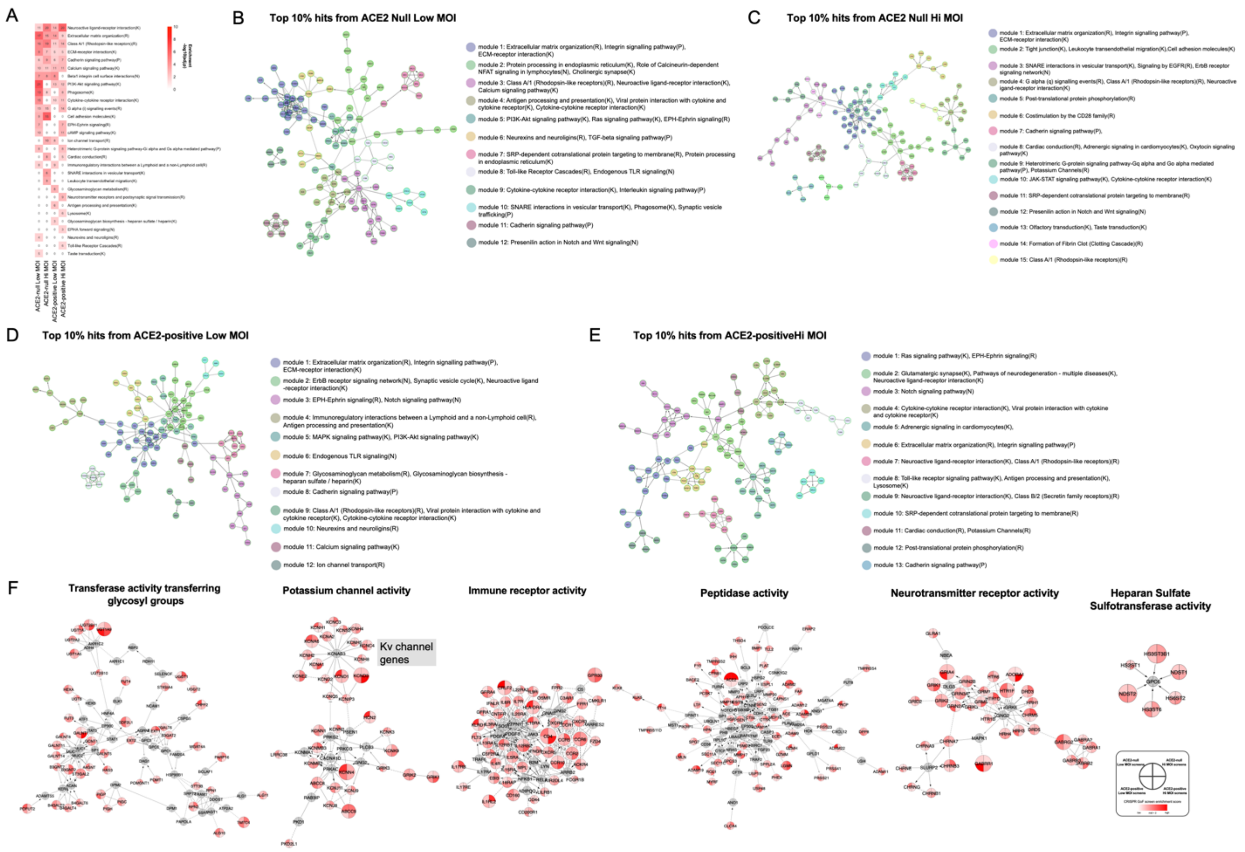


**Supplementary Fig. 4. Additional functional network analysis of the top hits in our screen.** (a) Heatmap shows the gene sets enrichment in different screen conditions. (b-e) Functional network clusters using the top 10% of the hits in different screen conditions. (f) Full functional network clusters involved with the top GO terms, refer to **Fig. 2d**.

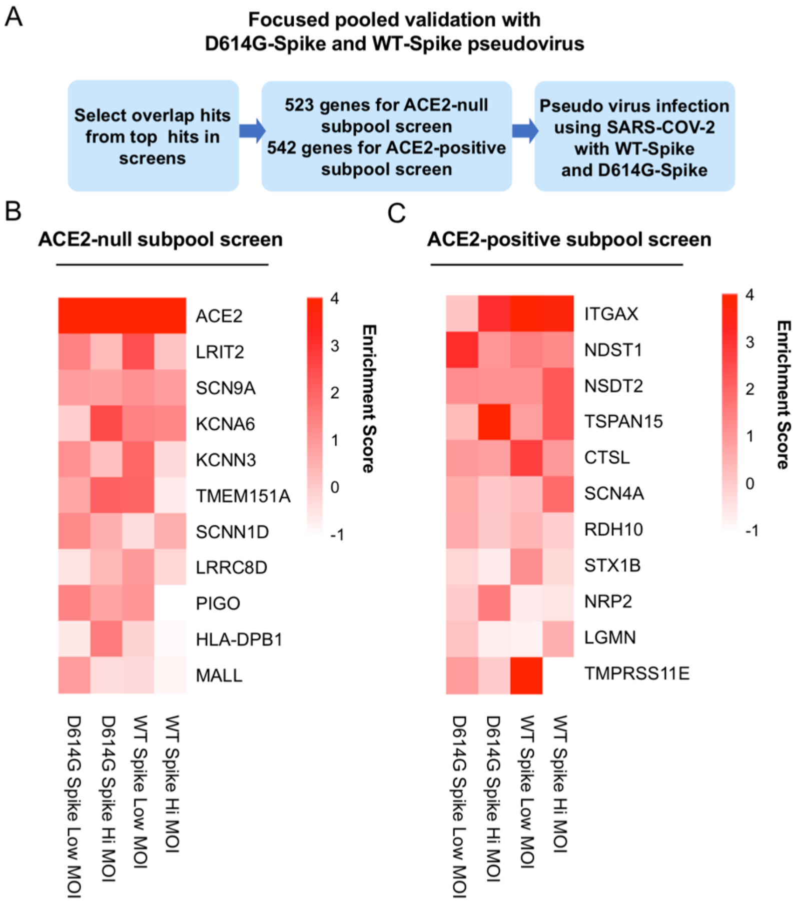


**Supplementary Fig. 5. Focused pooled validation using SARS-CoV-2 614G/641D Spike pseudotyped lentivirus.** (a) Workflow of the focused CRISPRa pooled validation screen. (b-c) Heatmaps show the enrichment scores of the top hits in (b) ACE2-null 293FT cells or (c) ACE2-positive 293FT cells.

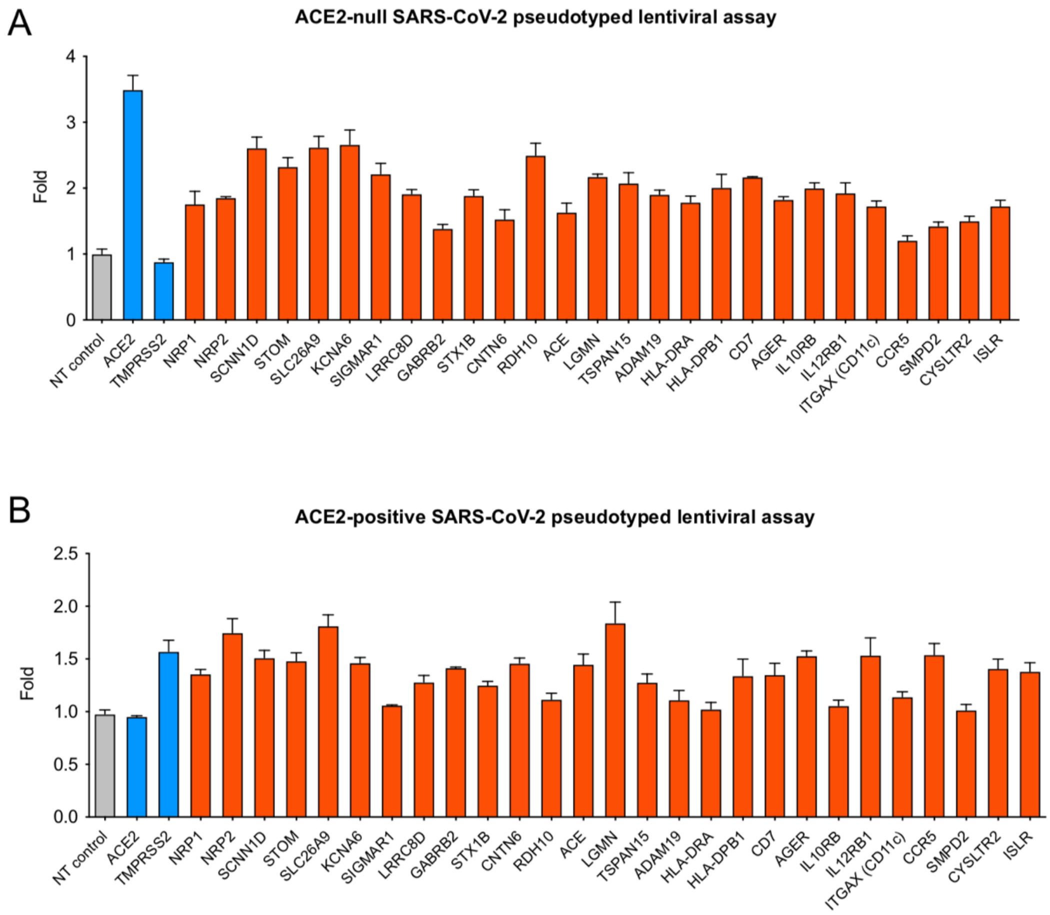


**Supplementary Fig. 6. Arrayed CRISPRa validation of screen hits using SARS-CoV-2 Spike pseudotyped lentiviruses.** SARS-CoV-2 Spike pseudotyped lentiviruses were used to transduce (a) ACE2-null or (b) ACE2-positive 293FT/dCas9-VP64 cells stably expressing different sgRNAs targeting the promoters of genes of interest.

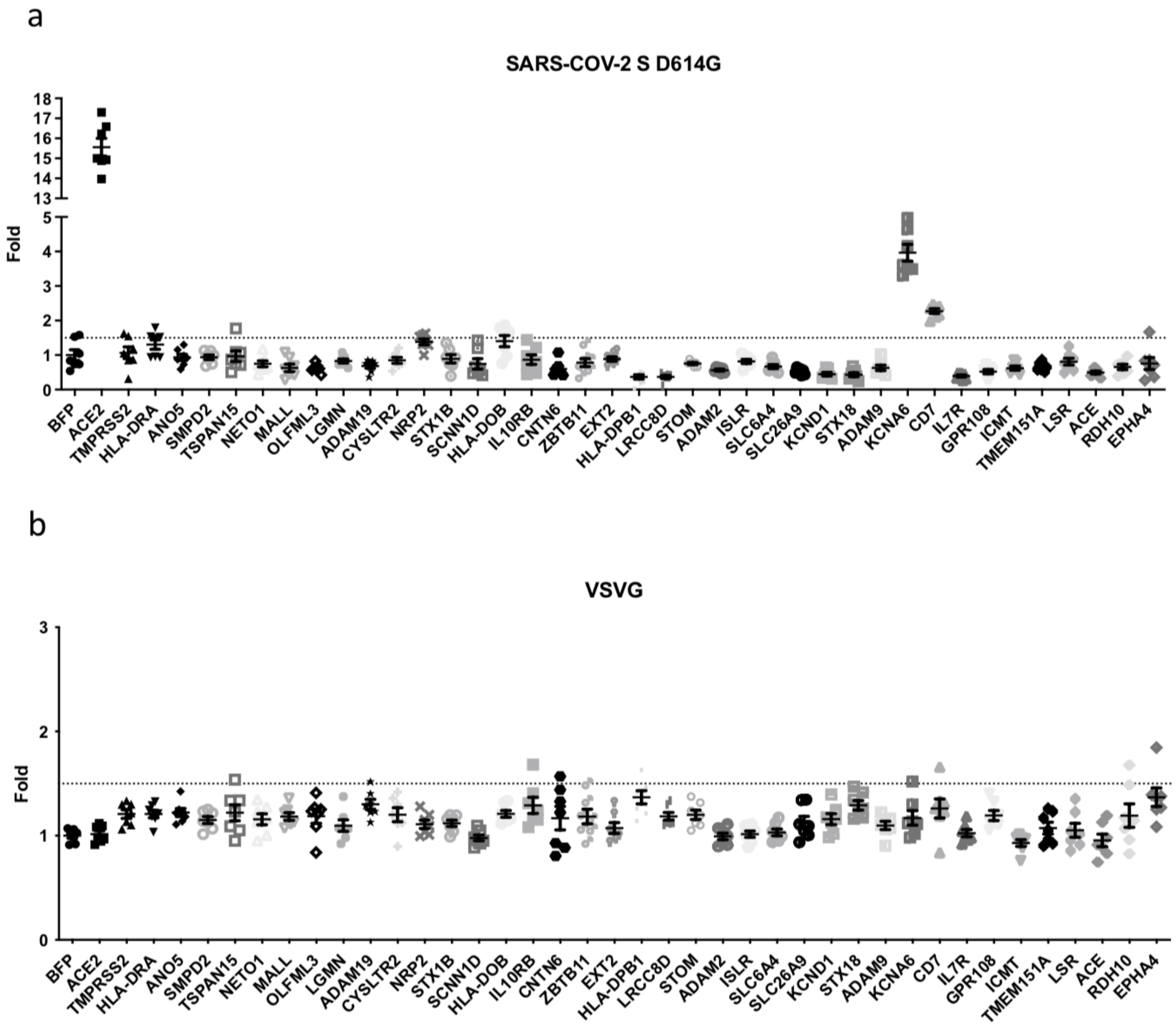


**Supplementary Fig. 7. Pseudotyped lentiviral assays in ACE2-null 293FT cells.** (a-b) ACE2-null lines stably overexpressing cDNAs of putative SARS-CoV-2 entry factors were transduced with lentiviruses pseudotyped with either (a) SARS-CoV-2 Spike D614G protein or (b) VSVG.

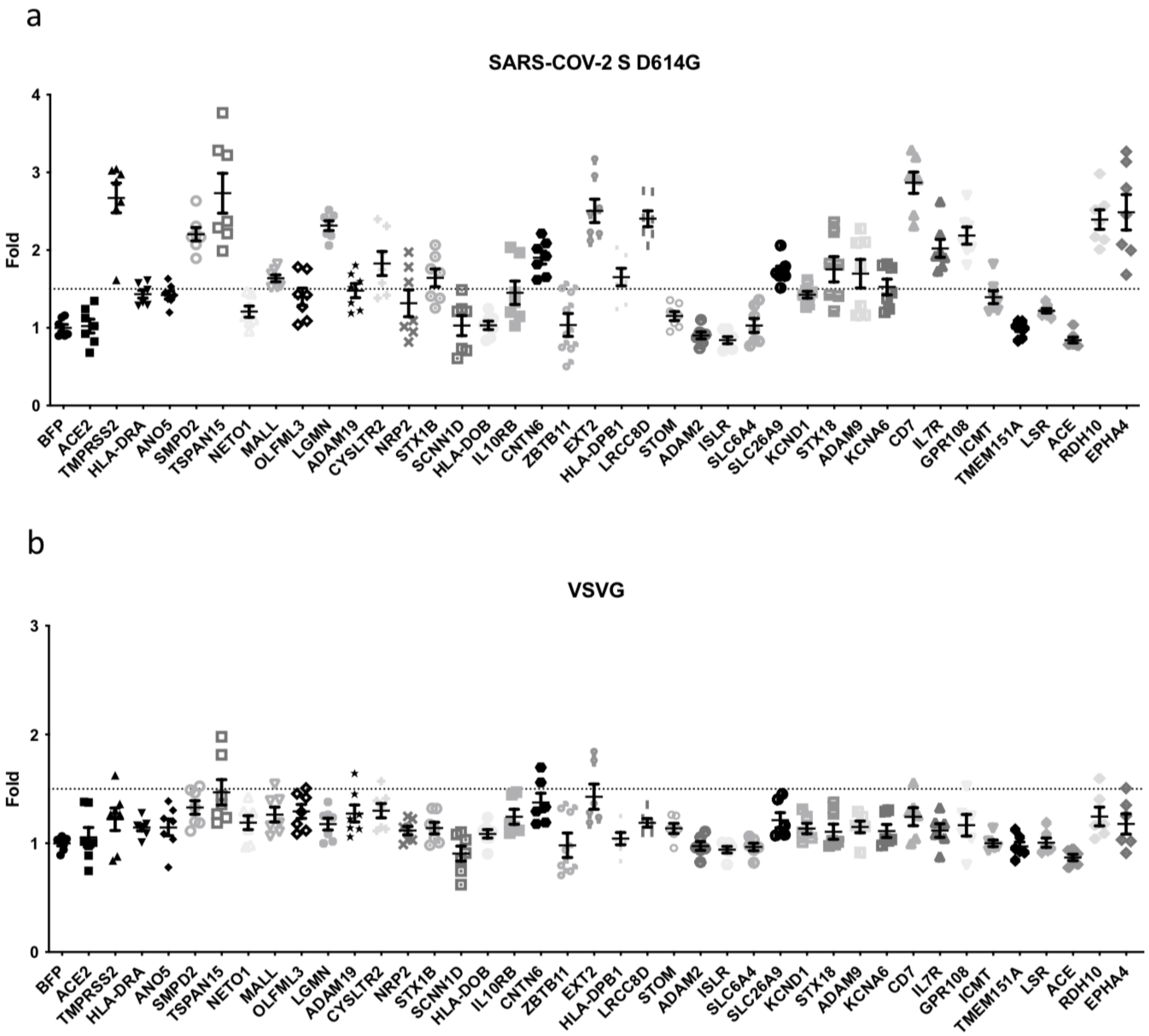


**Supplementary Fig. 8. Pseudotyped lentiviral assays in ACE2-positive 293FT cells.** (a-b) ACE2-positive lines stably overexpressing cDNAs of putative SARS-CoV-2 entry factors were transduced with lentiviruses pseudotyped with either (a) SARS-CoV-2 Spike D614G protein or (b) VSVG.

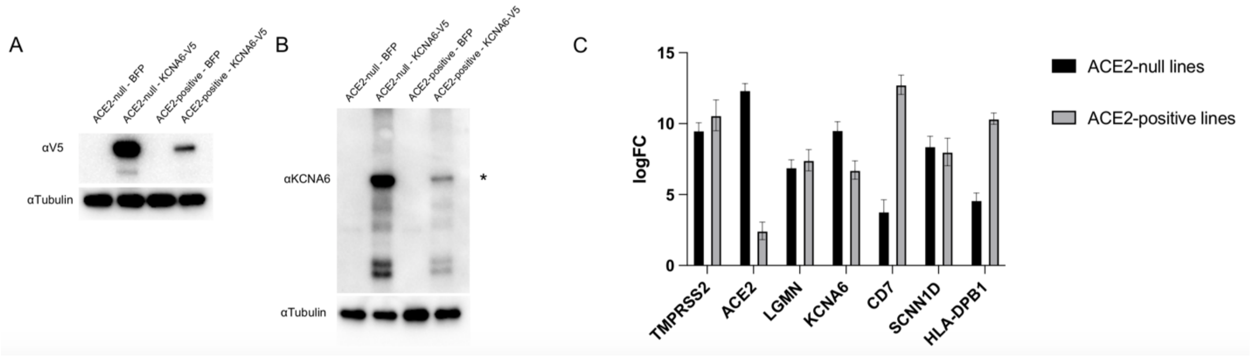


**Supplementary Fig. 9. Detection of cDNA expression in the overexpression cell lines.** (a-b) Western Blots of lysates from ACE2-null and ACE2-positive cells overexpressing either BFP or KCNA6 probed with either an (a) anti-V5 or (b) anti-KCNA6 antibody. * Denotes the correct band for KCNA6 based on protein size markers. (c) qPCR assay of the cDNA overexpressing cell lines. LogFC was calculated relative to BFP-overexpressing negative control cell lines.

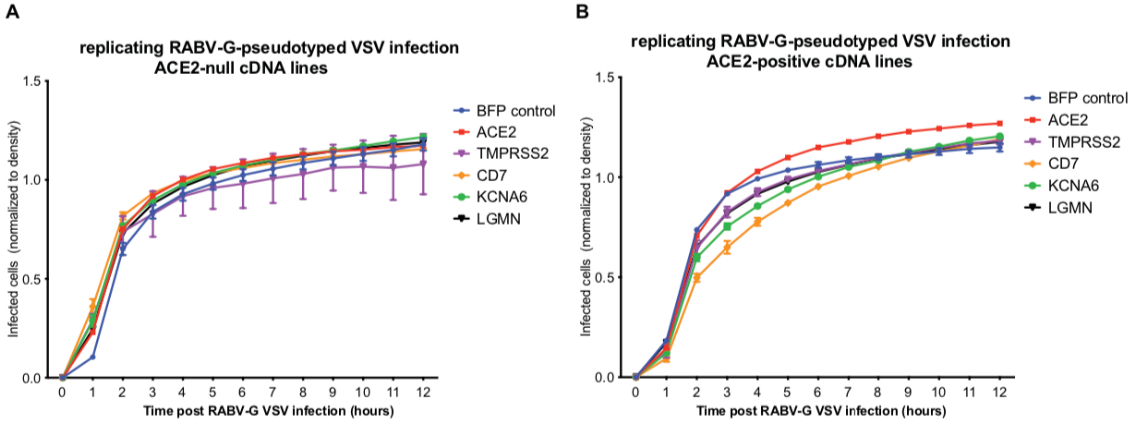


**Supplementary Fig. 10. Time-lapse imaging of replicating VSVdG-RABV-G SAD-B19 infection of cDNA overexpression lines.** (a-b) The levels of RABV-G pseudovirus infection tested in (a) ACE2-null and (b) ACE2-positive cDNA overexpression cell lines.

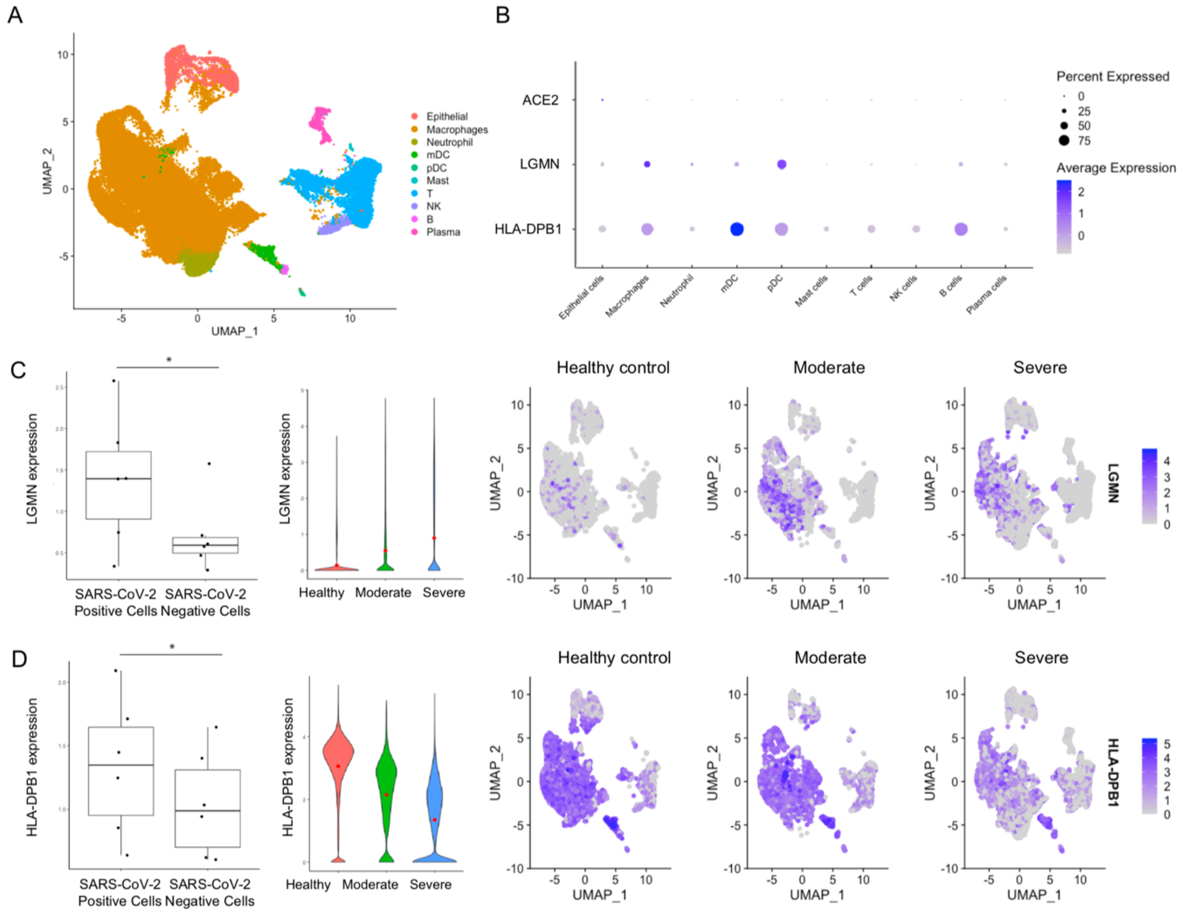


**Supplementary Fig. 11. RNA expression of LGMN and HLA-DPB1 in bronchoalveolar lavage fluids (BALF) of SARS-CoV-2 patients.** (a) UMAP depicting the major cell types and clusters in the BALF samples obtained from Liao *et al*. (n=13). (b) Dot plot visualization of the expression of ACE2, LGMN and HLA-DPB1 in BALF cells. (c-d) Left panels: boxplots comparing the average expression levels of LGMN and HLA-DPB1 between SARS-CoV-2 positive and negative cells from severely affected patients. Each dot represents a severely affected patient. Middle panels: LGMN and HLA-DPB1 expression of cells obtained from healthy controls, patients with moderate and severe COVID-19 infection. Right panels: UMAPs depicting the expression of LGMN and HLA-DPB1 in cells obtained from healthy controls (n=4) and COVID-19 patients (moderate, n=3; severe, n=6).

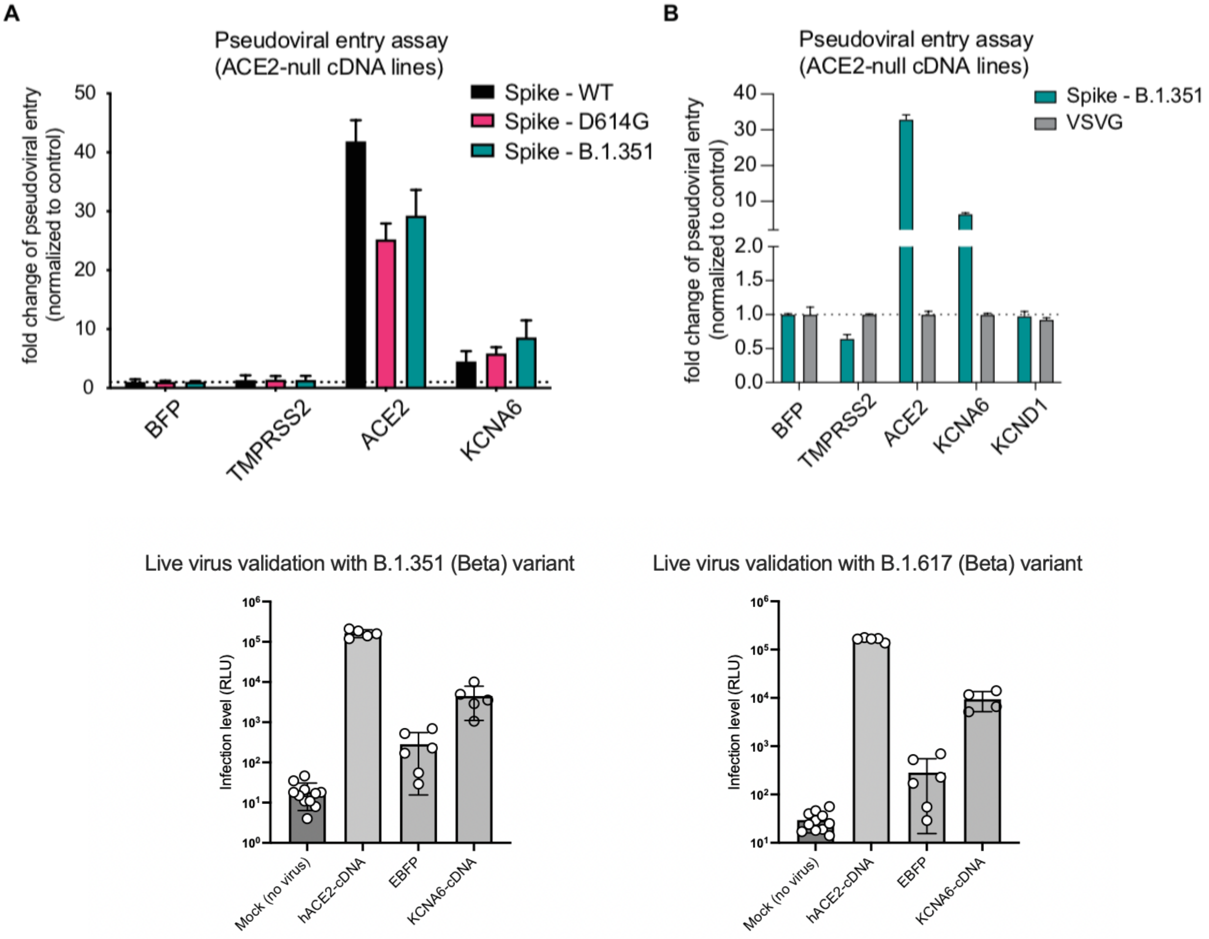


**Supplementary Fig. 12. Measuring pseudoviral entry using wild-type (WT) and mutated Spike variants (D614G and B.1.351) in cDNA overexpression lines.** (a) The levels of different Spike variants pseudovirus infection using ACE2-null cDNA overexpression cell lines. (b) Pseudoviral entry measured across different cDNA lines using Spike – B.1.351 variant and VSVG control. (c) Live virus validation of the effects of KCNA6 over SARS-CoV-2 infection using two different variants of concerns (Beta and Delta variants).
